# Scalable emulation of protein equilibrium ensembles with generative deep learning

**DOI:** 10.1101/2024.12.05.626885

**Authors:** Sarah Lewis, Tim Hempel, José Jiménez-Luna, Michael Gastegger, Yu Xie, Andrew Y. K. Foong, Victor García Satorras, Osama Abdin, Bastiaan S. Veeling, Iryna Zaporozhets, Yaoyi Chen, Soojung Yang, Arne Schneuing, Jigyasa Nigam, Federico Barbero, Vincent Stimper, Andrew Campbell, Jason Yim, Marten Lienen, Yu Shi, Shuxin Zheng, Hannes Schulz, Usman Munir, Ryota Tomioka, Cecilia Clementi, Frank Noé

**Affiliations:** AI for Science, Microsoft Research; Freie Universität Berlin, Department of Physics, Arnimallee 12, 14195 Berlin

## Abstract

Following the sequence and structure revolutions, predicting the dynamical mechanisms of proteins that implement biological function remains an outstanding scientific challenge. Several experimental techniques and molecular dynamics (MD) simulations can, in principle, determine conformational states, binding configurations and their probabilities, but suffer from low throughput. Here we develop a Biomolecular Emulator (BioEmu), a generative deep learning system that can generate thousands of statistically independent samples from the protein structure ensemble per hour on a single graphical processing unit. By leveraging novel training methods and vast data of protein structures, over 200 milliseconds of MD simulation, and experimental protein stabilities, BioEmu’s protein ensembles represent equilibrium in a range of challenging and practically relevant metrics. Qualitatively, BioEmu samples many functionally relevant conformational changes, ranging from formation of cryptic pockets, over unfolding of specific protein regions, to large-scale domain rearrangements. Quantitatively, BioEmu samples protein conformations with relative free energy errors around 1 kcal/mol, as validated against millisecond-timescale MD simulation and experimentally-measured protein stabilities. By simultaneously emulating structural ensembles and thermodynamic properties, BioEmu reveals mechanistic insights, such as the causes for fold destabilization of mutants, and can efficiently provide experimentally-testable hypotheses.

## 1 Introduction

Proteins and protein complexes constitute the functional building blocks of life and are at the center stage of drug development, enzymatic catalysis, biotechnological processes and biomaterials. Consequently, understanding how proteins work and how their function can be regulated or designed is one of the grand challenges in science and technology. Protein science can be characterized by three pillars of understanding: sequence, structure, and function. Next-generation sequencing has made it possible to acquire the protein sequences of entire genomes at low cost, while AlphaFold [1] and similar models [2–4] have built upon the decades of data accumulated in the Protein Data Bank (PDB) [5] to predict 3D protein structures that in many cases match experimental accuracy within minutes. For protein function, unfortunately, methods that are both highly accurate and high-throughput are missing, and thus our understanding of how proteins work remains anecdotal.

Functional descriptions such as “actin builds up muscle fibers” are human-made attributions that arise from objectively measurable mechanistic properties: (i) What are the conformational states (i.e., sets of different structures) a protein can be in? (ii) Which other molecules can a protein bind to in these different conformations? (iii) What is the probability of these conformational and binding states at a given set of experimental conditions? For example, actin exists in multiple conformational and binding states that are regulated by its cofactors ATP/ADP (Fig. 1a), providing the molecular basis of muscle growth.

**Fig. 1.**
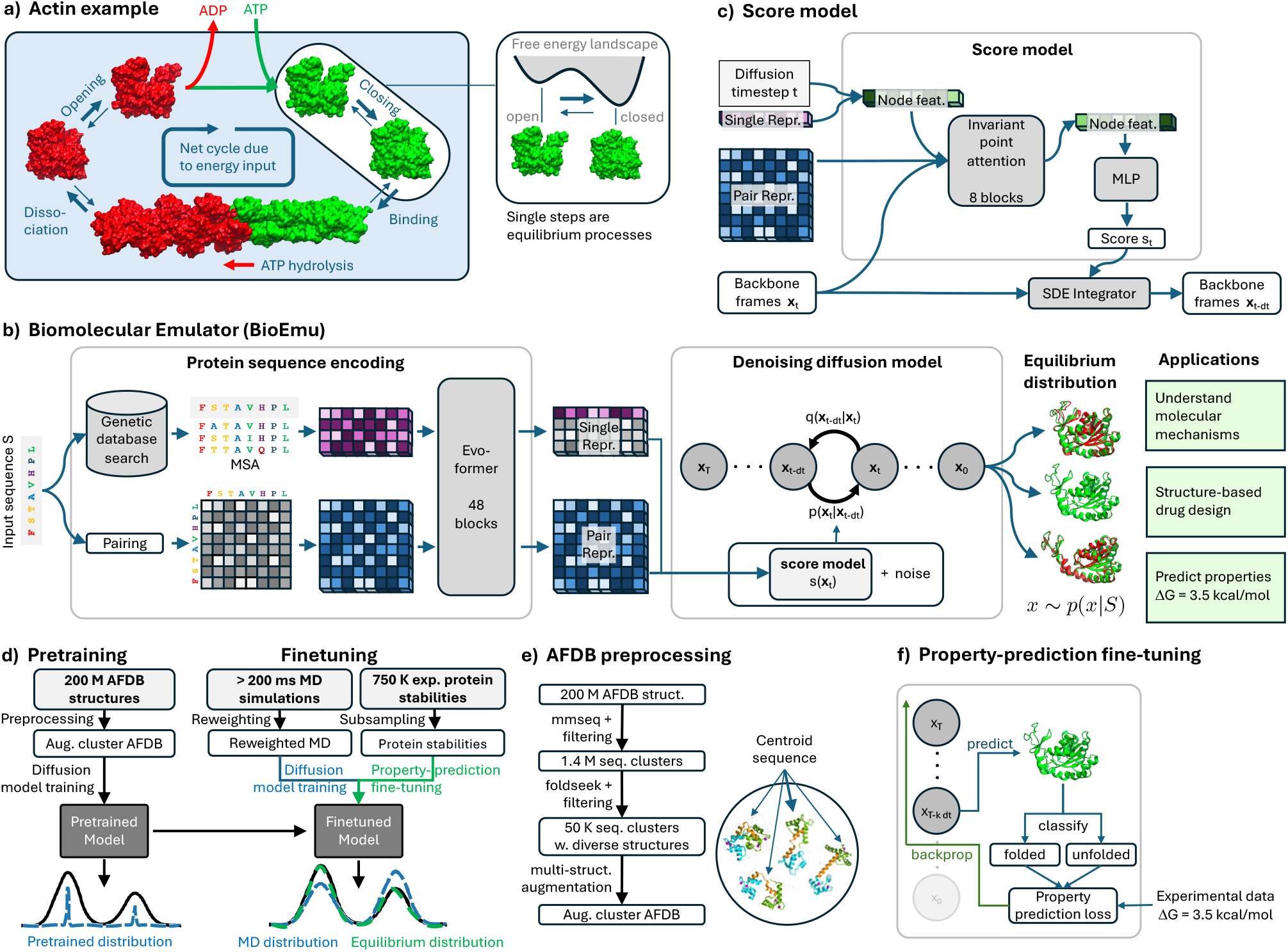
Overview of model and architecture. a) Actin conformational changes and filament formation / dissociation as an example for the mechanistic basis of protein function. b) ML model architecture consisting of protein sequence encoder and denoising diffusion model. The diffusion model samples coarse-grained protein structures from an approximate equilibrium distribution, from which properties such as free energy differences can be computed. c) Architecture of the score model used in the denoising diffusion model. d) Data integration and model training pipeline. e) Data processing pipeline for pretraining. f) Experimental property training for finetuning.

Available technologies that probe such conformational and binding states and their probabilities at high accuracy are currently not scalable. Single-molecule experiments can provide the full equilibrium distributions of observables such as intramolecular distances [6], but require bespoke molecular constructs and time-consuming data collection. Cryo-electron microscopy can resolve multiple conformational states of biomolecular complexes along with their probabilities [7], but running these experiments is costly both from a monetary and time perspective. Molecular Dynamics (MD) simulation is, in principle, a universal tool that allows both structure and dynamics of biomolecules to be explored at all-atom resolution. However, biomolecular forcefields are far from perfect and the sampling problem renders the study of protein folding or association via MD a feat of epic computational costs for small-sized proteins, even if special-purpose supercomputers or enhanced sampling methods are employed [8, 9]. Machine-learned coarse-grained MD models have an opportunity to achieve similar accuracy as all-atom MD at 2-3 orders of magnitude lower computational cost [10, 11] but are still under development.

The grand challenge to complete our understanding of protein function thus motivates the development of a technology that can help elucidate protein conformational states and binding states, as well as their associated probabilities. This technology should ideally achieve an accuracy comparable to a converged MD simulation, or a cryo-EM experiment with multi-conformation analysis, but it should only require a few hours of wall-clock time and cost no more than a few dollars per experiment. Generative systems, such as Boltzmann Generators [12] (BGs), which can efficiently sample arbitrarily-defined equilibrium distributions, indicate that such technologies may be within reach, but are difficult to scale to large proteins. Concurrently, diffusion models and similar approaches are now widely used in protein structure prediction and design [2, 3]. Such models [13–15], as well as perturbation-based derivatives of AlphaFold [16, 17] have also been shown to be capable of generating distinct protein structures and can be combined with MD simulation to alleviate the sampling problem [18]. As yet, generative ML systems have mainly demonstrated an ability to qualitatively sample distinct protein conformational states. A demonstration that generative ML can quantitatively match equilibrium ensembles and predict experimental observables is critical going forward [19].

Here we set out to develop a first version of an ML system that can approximately sample from the equilibrium distribution of protein conformations within a few GPU-hours per experiment — a **biomolecular emulator (BioEmu)**. The biggest challenge in training such a generative model is that no single high-quality data source for training exists due to the aforementioned challenges with experimental methods and MD. We therefore train BioEmu by combining data ranging from a large set of static protein structures and vast amounts of MD simulation to experimental measurements of protein stabilities. We validate the system on a range of tasks: (i) the prediction of protein conformational changes including large domain motions, local unfolding, and the formation of cryptic binding pockets, (ii) the emulation of equilibrium distributions that can be generated by high-throughput MD simulation, and (iii) the prediction of experimentally-measured stabilities of folded states of small proteins by directly generating equilibrium ensembles and explaining structure-stability relationships of mutants. We demonstrate that free energies can be predicted with errors below 1 kcal/mol and are therefore on the order of experimental accuracy.

Given its versatility and efficiency, we believe that BioEmu has a variety of practical use cases, ranging from helping with current MD simulation workflows, the interpretation of protein experiments, identification of binding pockets and allosteric mechanisms in drug discovery, and generation of ensembles for dynamical protein design. Importantly, our demonstration that the large upfront costs of MD simulation and experimental data generation can be amortized and the prediction error decreases with an increasing amount of diverse training data indicates a path forward for predicting biomolecular function at genomic scale.

## 2 Model

BioEmu uses a similar model architecture as Distributional Graphormer [13], but with a significantly different training approach. Starting from the input protein sequence, single and pair representations of the sequence are computed using the AlphaFold2 evoformer [1]. These sequence representations serve as input to a denoising diffusion model that generates protein structures (Fig. 1b,c; Sec. S.2). Sequence encoding is invoked only once per protein, and using a second-order integration scheme we generate protein structures in as few as 100 denoising steps (Sec. S.2.3), leading to high sampling efficiency: 10,000 independent protein structures from the learned equilibrium distribution can be sampled within minutes to a few hours on a single GPU, depending on their size.

For model training and testing we have developed several new benchmarks and training methods to integrate the heterogeneous data modalities (Sec. S.1, S.3). BioEmu is pretrained on a clustered version of the AlphaFold database (AFDB), using a data augmentation strategy that incentivizes it to sample diverse conformations (Fig. 1d,e, Sec. S.3.2). Starting from this pretrained model, we then continue to train on a mixture of MD data and experimental measurements of protein stability, plus occasional examples from the pretraining data. We have curated and generated a total of over 200 milliseconds of all-atom MD data for small-to-medium proteins (Sec. S.3). To mitigate the sampling problem, MD data was reweighed towards equilibrium using either Markov State Models [20], or weights from experimental data (see S.3.5.3), when possible. The reweighed MD data is used in a second training stage of the model (Fig. 1d). The experimental measurements of protein stability that we train on are a subset of the MEGAscale dataset [21], which comprises on the order of a million protein stability measurements (Fig. 1d). As the MEGAscale dataset does not contain structures, we developed an new algorithm called property-prediction fine-tuning (PPFT) to efficiently incorporate experimental measurements into diffusion model training (Fig. 1f, Sec. S.3.6). Finally, to evaluate generalization, we filter our training set such that no protein has more than 40% sequence similarity to any of the reported test proteins of at least 20 residues or longer. The model name BioEmu denotes the fine-tuned model, trained on AFDB, MD simulations and experimental measurements of protein stability. Subsequent results use this model unless otherwise described.

## 3 Sampling conformational changes related to protein function

We regard the ability to sample distinct biologically relevant conformations qualitatively as a basis to build a quantitative equilibrium sampler. Therefore we first test qualitatively if BioEmu’s samples include known conformational changes and compare this capability with AFCluster [16] and AlphaFlow [14] as two representative baseline methods. Towards this goal, we defined a challenging test set of conformational changes, called OOD60, with a maximum of 60% and 40% sequence similarity to the AlphaFold2 monomer model and our training sets, respectively. Due to the strict sequence similarity constraints, OOD60 only contains 19 proteins, but it features various challenging cases like large-scale conformational changes caused by binding to other biomolecules (Fig. S1). While it is uncertain if all of these conformational changes can be predicted by a single-domain model, the benchmark tests for strong generalization and we find that our model significantly outperforms the two considered baseline approaches (Fig. S5a).

In order to evaluate the multi-conformation capabilities of our model more exhaustively, we have also curated a set of around 100 proteins that engage in experimentally-validated domain motions, local unfolding transitions, or cryptic pocket formation. These include some proteins contained in OOD60 as well as proteins that overlap with the AlphaFold2 training set. We confirmed that the model’s performance is similar for proteins that overlap with the AlphaFold2 training set and those that do not, indicating that the benchmark does not test capabilities that the model trivially extracted from evoformer embeddings (Table S4). Furthermore, BioEmu outperforms other methods except for the apo states in the cryptic pocket benchmark, and the difference is especially large for the proteins outside the AlphaFold2 training set (Fig. S5, b-d).

Our curated benchmark furthermore demonstrates that our model qualitatively captures functionally relevant protein conformations. For example, proteins can undergo large-scale domain motions as part of their functional cycle. In the open-close transition of Adenylate Kinase, the closed state brings the substrates together to catalyze the ATP + AMP ⇌ 2ADP reaction. Single-molecule experiments have confirmed that opening and closing occurs reversibly on timescales of tens of microseconds when the substrates are bound [22]. BioEmu predicts a range of open and closed states, including close matches with crystallographic structures (Fig. 2a,i). A second example is the open-close transition of LAO-binding protein which is required to bind and release lysine, arginine and ornithine for transport across membranes as part of the ATP-binding cassette protein family (Fig. 2a,ii). Another interesting example of domain motions is that of the receptor module which regulates the concentration of cyclic di-GMP in bacteria. In this case one domain undergoes a large-scale rotation and repacks to the other domain with a completely different contact pattern (Fig. 2a, iii). See Fig. S2 for 15 further examples. Overall, BioEmu predicts 85% of the reference experimental structures with ≤3 Å RMSD (Fig. 2a), indicating the model’s ability to predict which protein regions are more or less flexible, as well as which resulting motions can occur.

**Fig. 2.**
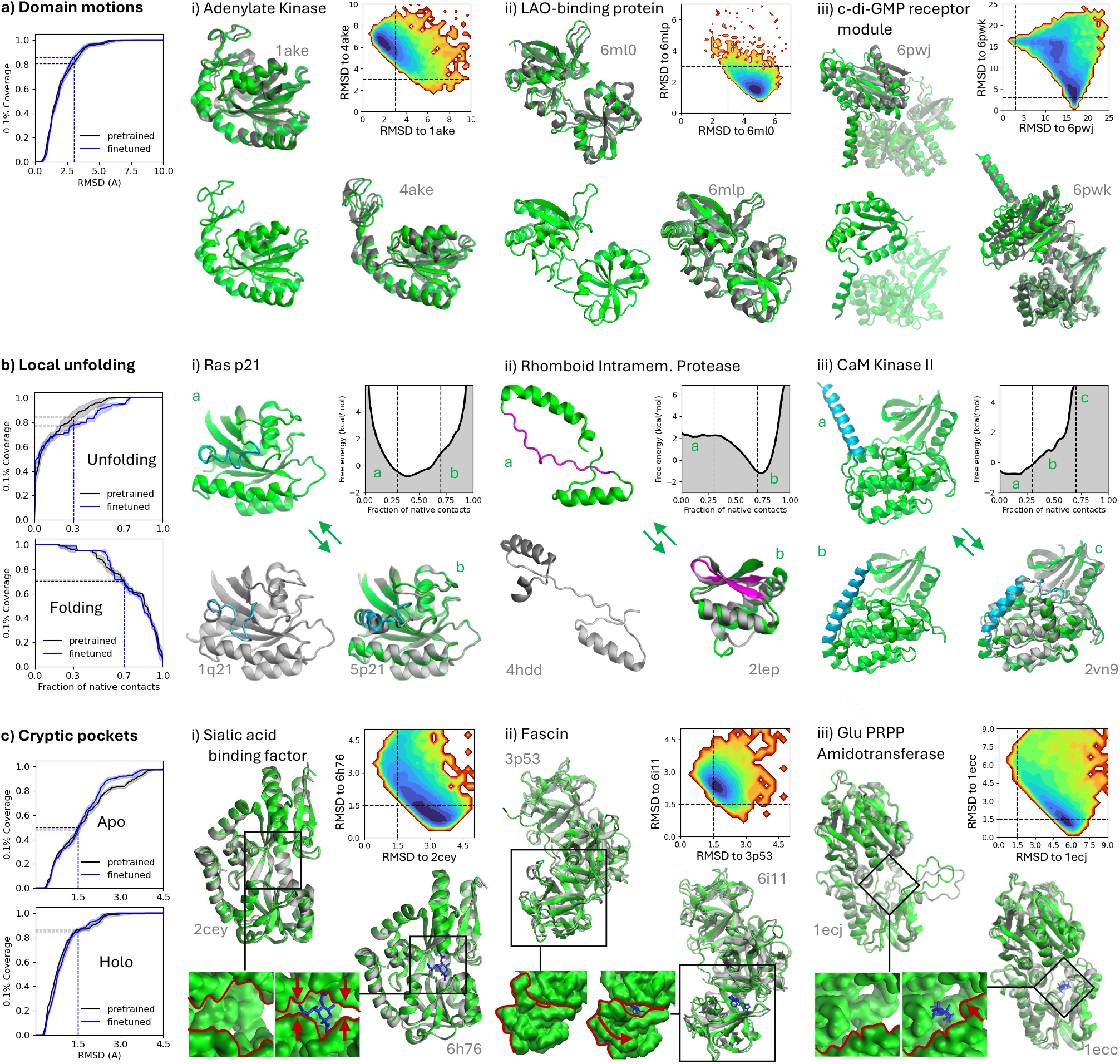
BioEmu samples functionally distinct protein conformations. a) Large-scale domain motions such as opening/closing, rotation, and repacking. b) Local unfolding or unbinding of parts of the protein. c) Formation of cryptic binding pockets that are not present in the apo ground state. Left column shows coverage, defined as the percentage of reference structures that are sampled by at least 0.1% of samples (4 kcal/mol) within a given distance of the respective metric. ‘Pretrained’: results after training on only AFDB data; ‘finetuned’: results using BioEmu. Global and local root mean square deviation (RMSD) is used for domain motions and cryptic pocket formation benchmarks, respectively, and fraction of native contacts (FNC) for local unfolding. Our defined success threshold is marked by dashed lines. i), ii), iii) show three examples for each class of evaluated conformational changes, using BioEmu. All examples shown have less than 40% sequence similarity to BioEmu’s training data. Six references are also not included in the AlphaFold2 monomer model training set (both in a,ii-iii both references and holo states in c,i-ii). The example shown in a,iii is also in the OOD60 benchmark set, having less than 60% sequence similarity to any protein in the AlphaFold2 training set.

Next we consider local unfolding transitions, in which part of a protein chain unfolds or detaches from its main structure as part of a signaling pathway. Predicting local unfolding is arguably more challenging than predicting domain motions, as it requires the model to correctly rank which parts of a protein’s fold are more stable. A famous example of local unfolding is Ras p21, a conformational switch which signals cell growth and whose mutants are often linked to cancer development [23] (Fig. 2b,i). In its active state, stabilized by GDP binding, the Switch II region forms a short alpha-helix, which partially unfolds in the inactive state stabilized by GTP. Rhomboid intramembrane protease (Fig. 2b,ii) is a much more complex case of domain swapping. Its monomeric form features a globular conformation, while in its dimeric form the central beta-sheet unfolds and the helices of the two monomers bind to each other. Finally, CaM Kinase II (Fig. 2b,iii) presents an autoinhibition mechanism, in which the N-terminus binds into the active site. BioEmu predicts these local unfolding transitions correctly, and overall 72% of locally folded and 74% of locally unfolded states across 20 protein examples (Fig. 2b, Fig. S2).

As a third class of conformational changes we consider the formation of pockets that are absent in the apo state but form upon small-molecule binding. One option to identify such “cryptic” binding pockets is high-performance MD simulation [24], but the millisecond timescales often involved in the spontaneous opening of such pockets make MD on commercial hardware rarely viable for in-silico drug discovery pipelines. We have curated 34 cases of experimentally-validated formation of cryptic binding pockets from the literature (Fig. S4). The sialic acid binding factor (Fig. 2c,i) presents a case where a large opening in the apo state can partially close and form a binding site for the ligand. Fascin is a four-domain protein where two domains can rotate with respect to each other, to reveal a binding site (Fig. 2c,ii). In Glu PRPP amidotransferase, part of the chain is unfolded in the apo site and folds into a structure that completes the binding site for the ligand (Fig. 2c,iii). To ensure capturing subtle changes, we define success by a very strict 1.5Å RMSD threshold to the apo and holo reference structures. Surprisingly the model has a strong preference for holo states and successfully predicts the cryptic pocket in 85% of cases, while it only succeeds in predicting 49% of the apo structures, indicating further room for improvement.

## 4 Emulating MD equilibrium distributions

A major motivation for the development of BioEmu is to side-step the infamous MD sampling problem. The practical MD data requirements for discovering biomolecular conformations and estimating their free energy differences are often in the range of 100 *μ*s to 10 *m*s simulation time [8, 9, 26]. Due to the vast computational costs, exhaustive MD sampling of biomolecules has only been achieved in few cases, either with special purpose supercomputers [27] or via large-scale distributed simulations integrated via statistical models [9, 28]. Here, we assess BioEmu’s ability to emulate the equilibrium distribution that would be sampled with extensive MD simulations. To this end, we have amassed all-atom simulations of proteins with a total aggregated simulation time of over 200 ms (Table S1), which are used for fine-tuning BioEmu (Fig. 1d).

Before analyzing the model trained on the full dataset, we first test whether BioEmu’s design permits learning to emulate long-time MD equilibrium distributions, using D. E. Shaw research (DESRES) simulations of 12 fast-folding proteins generated on the special-purpose supercomputer Anton [8]. We train 12 “DESRES-finetuned models”, each of which is tested on one fast folder and fine-tuned on the others. As expected, the AFDB-pretrained model predicts the native state but exhibits poor performance in free energy surface sampling (Fig. S7). However, fine-tuning on only 11 sequences results in a surprisingly good match in the free energy surfaces of test proteins (Fig. 3a,i, Fig. S7). For all proteins, the model predicts both native as well as the unfolded states with similar shapes on the free energy landscape. In many cases, several or all folding intermediates visible on a two-dimensional free energy surface are predicted (Fig. 3a,i, Fig. S7): For beta-beta-alpha protein (BBA), both MD and the DESRES-finetuned models predict the existence of an intermediate with the alpha-helix formed and the beta-sheet broken. For protein G, both MD and the DESRES-finetuned models sample intermediates with half of the beta-sheet still formed, while the other half and most of the helix are broken. For homeodomain, MD and model agree in the prediction of an intermediate with only one helix turn unwound, while the unfolded states still feature some degree of helical propensity. There is an excellent agreement of the predicted secondary structure propensities with the MD data (Fig. 3a, i, rightmost column). Quantitatively, the mean average error between the MD and model 2D free energy landscapes is only 0.74 kcal/mol, ranging from 0.30 kcal/mol for BBA to 1.63 kcal/mol for *λ*-repressor, which is on the order of differences expected from two different classical MD force fields [29, 30].

**Fig. 3.**
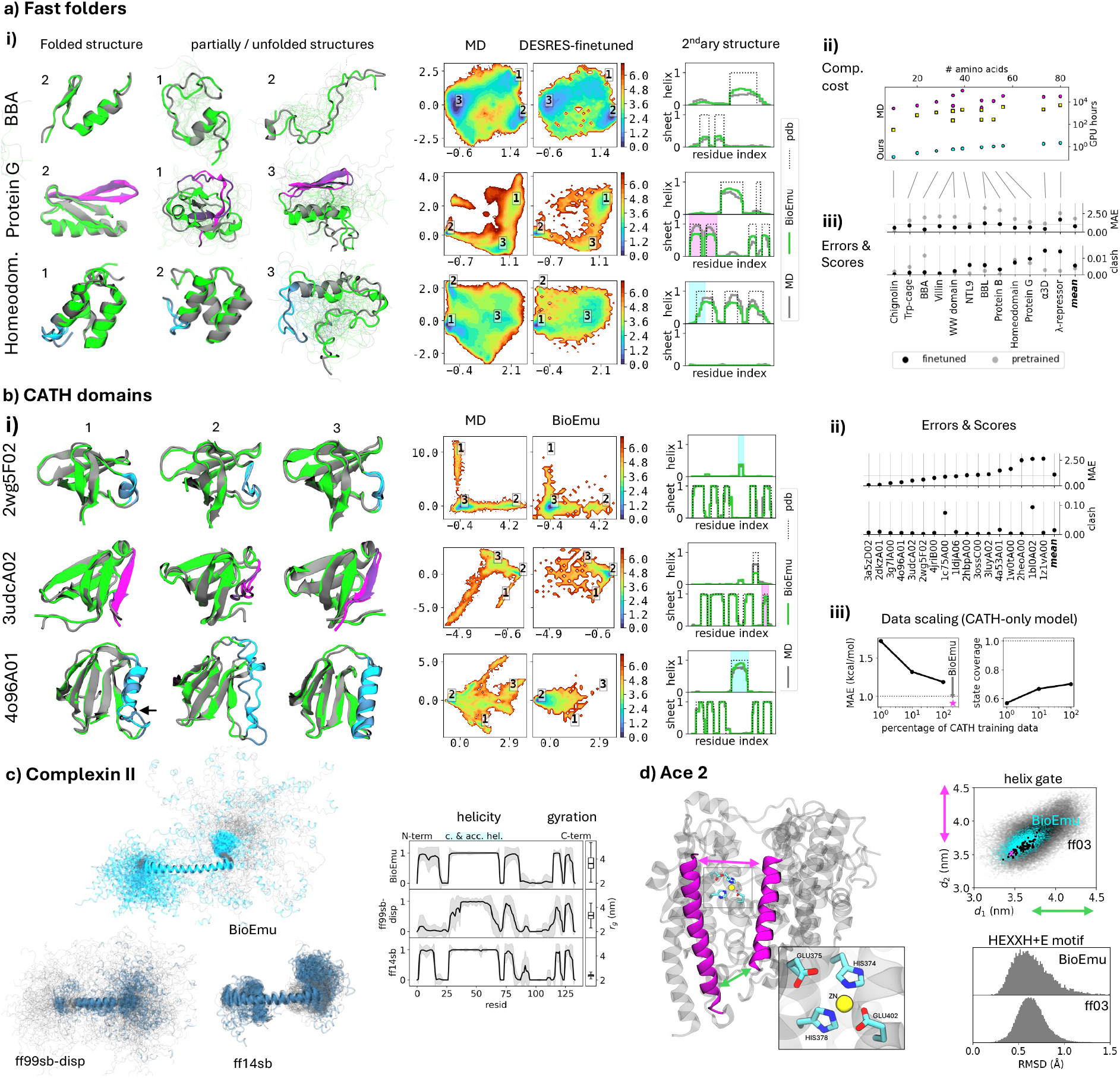
BioEmu emulates equilibrium distributions of all-atom molecular dynamics (MD) many orders of magnitude faster. a) Fast-folding proteins simulated by the DESRES Anton supercomputer compared with a model fine-tuned on all DESRES fast-folder except the test protein. i) From left to right: Folded and partially / unfolded structures predicted by our model (green) and ground truth MD (grey). Free energy surfaces (in kcal/mol) of ground truth MD and our model in the space of the two slowest time-laged independent components (TICA) [25]. Secondary structure content is compared over the whole ensemble of structures. ii) Computational cost (in GPU hours) for MD (magenta: full DESRES dataset; yellow: single folding-unfolding roundtrip) and 10k samples from our model (cyan). iii) mean average error (MAE) of free energy differences of macrostates and fraction of unphysical model samples due to clashes. b) CATH domains. i) Color-code as in a,i. Additionally, structurally flexible motifs are color-coded (cyan: helical; magenta: sheet) with reference MD-structures in dark and BioEmu samples in light color. ii) as in a, iii. iii) Macrostate free energy MAE and state coverage as function of training data of a specialized CATH-only model. MAE of BioEmu is indicated by a magenta star. c) Intrinsically disordered protein Complexin II: Sampled structure ensemble (left), helix content and radius of gyration (right) compared betweenBioEmu and two all-atom force fields. d) Conformational flexibility of human angiotensin-converting enzyme 2 (ACE2). Helix gate opening distribution as a function of two distances (magenta and green arrows) of extensive MD simulation (grey, seeded from magenta PDB-ID 6LZG), homologous PDB structures (black squares) and BioEmu samples (cyan). Backbone-RMSD to crystal structure of HEXXH+E motif compared between MD and BioEmu.

We compare the computational costs between MD data generation and BioEmu in GPU-hours (here on a NVIDIA Titan V). For all BioEmu results shown here, we draw 10k samples, which incurs computational costs of *<* 1 GPU-minute for Chignolin to around 1 GPU-hour for *λ*-repressor (Fig. 3a). For MD we consider the cost for generating the DESRES simulations, whose lengths have been chosen to include roughly 10 folding-unfolding transitions (Fig. 3a). The MD costs then range from 2,000 GPU-hours for Chignolin to more than 100,000 GPU-hours for NTL9, resulting in a model speedup advantage over MD of four to five orders of magnitude. We also note that for most proteins shown here, performing sufficiently long MD simulations to directly observe folding and unfolding in single trajectories is still not possible on consumer-grade hardware but instead requires a much more complex methodological framework [28, 31].

The main BioEmu model is fine-tuned on more than 200 ms MD simulations with Amber force fields at or near a temperature of 300K (Table S1). We choose to combine data from slightly different simulation conditions as each of these MD models is inherently imperfect, and we regarded experimental data as being more reliable for weighing between conformations (Fig. 1d, Sec. 5). Differences in the simulation conditions of our own generated data are intentional, e.g. AMBER ff99sb-disp [32] was chosen to avoid spuriously-misfolded states produced by other force fields in the context of protein folding (Sec. S.1.5.4). A large fraction of training data, 46 ms, is dedicated to 1100 CATH domains, common building blocks of protein structure [33] (Sec. S.1.5.2, S.1.5.3). We designate 17 CATH systems with more than 100 *μ*s simulation time as test set and report statistics comparing MD and model distributions (Fig. 3b, Fig. S8). Similar as for DESRES simulations, BioEmu predicts the native state with local fluctuations and typically several other substrates and structures sampled by MD. Most secondary structure propensities match well (Fig. 3b, rightmost column). We observed a free energy mean average error over the converged test set of 0.91 kcal/mol, again comparable to the differences expected between different MD force fields.

To understand whether our model’s ability to sample accurate equilibrium distributions is limited by training data or model expressivity, we trained 3 models with the same architecture as BioEmu from scratch, using only CATH data. We fixed a test set of CATH domains and trained the three models using 1%, 10% and 100% respectively of the remaining CATH domains. We observed decreased free energy errors and an increased coverage of the conformations sampled by MD as the amount of training proteins increased (Fig. 3b,iii), suggesting that the model can be further improved by adding more training data.

Finally, we have evaluated BioEmu for two case studies that involve larger proteins: Complexin II (134 aa) and ACE2 (614 aa). Complexin II is an intrinsically disordered protein (IDP) from the neurotransmitter release apparatus [34]. IDPs tend to be difficult to sample with MD, however, BioEmu can efficiently emulate a flexible ensemble of complexin II structures (Fig. 3c) while reproducing known secondary structure elements such as the central and accessory helices [34, 35]. Achieving convergence of IDPs of this size with all-atom MD is unpractical. At an orders of magnitude higher computational cost than with BioEmu, we have conducted ~5*μs* of MD simulations with all-atom MD, which are most likely not converged but already display qualitatively different behavior: The AMBER ff14sb force field produces a very rigid compact structure with a small radius of gyration and little to no variation in secondary structure content, whereas AMBER ff99sb-disp tends to destabilize known secondary structure elements (Fig. 3c). The second case-study is ACE2, a metalloenzyme that plays an important role in SARS-CoV-2 viral uptake (Fig. 3d) [36]. We show that BioEmu reliably reproduces a stable HEXXH+E motif, the active enzymatic site of this protein [36], with backbone RMSDs below 1.5Å for all model samples. The distance distribution of two gate-keeping helices is similar albeit less flexible compared to a vast set of molecular dynamics simulations (890 *μ*s, Ref. [37]) and compatible to homologue PDB structures (Fig. 3d).

## 5 Predicting protein stabilities

Understanding protein stability is crucial for various applications in molecular biology, drug design, and biotechnology. From a modeling point of view, predicting a protein’s stability is a specific case of predicting the equilibrium probabilities of its different conformational states, and these all arise from the same underlying biophysics. We therefore desire to train BioEmu so that the proportion of samples in folded and unfolded states matches the experimentally-measured protein stability. We measure prediction errors in terms of the folding free energy, defined as Δ*G* = *G*_folded_ − *G*_unfolded_, and classify protein structures as folded or unfolded based on their fraction of native contacts (Sec. S.3.5.3).

To facilitate protein stability prediction, BioEmu’s training data includes over 750,000 experimental measurements from the MEGAscale dataset (Sec. S.1.7) [21], with a total of 25 ms of all-atom MD simulations of the folded and unfolded states of 271 wildtype proteins and 21458 mutants (S.1.5.4). To address MD sampling and force field issues, we weigh the folded and unfolded samples so that they correspond to the experimentally-measured protein stabilities (Fig. 1d). To speed up training convergence and leverage the large number of MEGAscale measurements, we developed the Property Prediction Fine-Tuning algorithm (PPFT, Fig. 1f, S.3.6) that integrates experimental expectation values, such as protein stabilities, into diffusion model training without requiring protein structures. PPFT uses fast approximate sampling with only 8 denoising steps, which we observed to be sufficient to confidently predict whether each sampled structure will be classified as folded or unfolded. By comparing the mean foldedness of sampled structures with experimental measurements and backpropagating the error, our model can be efficiently trained to match experimental protein stabilities.

Our model achieves a mean absolute error below 0.8 kcal/mol and a Spearman correlation coefficient above 0.65 for proteins in the MEGAscale dataset (Fig. 4a). This accuracy outperforms other existing black box methods that predict Δ*G* values directly from sequences [38–41]. Interestingly, the errors are similar for both training and test set, indicating that the model generalizes well but cannot perfectly fit the training data, perhaps due to inconsistencies between the folded/unfolded state definitions between this work and what the experiment is sensitive to. To check whether BioEmu makes physically reasonable predictions outside the MEGAscale set of proteins, we tested it on proteins that are known to be both very stable and unstable. We first selected stable proteins from ProThermDB [42] with Δ*G <* −8 kcal/mol (more details in S.5.1). Our model consistently samples these proteins in their folded states with a fraction of native contacts always exceeding the 0.65 threshold. To test whether our model systematically predicts intrinsically disordered proteins (IDPs) as unfolded, we used the CALVADOS test set [43]. Most proteins sampled displayed a radii of gyration (*R*_*g*_) similar to random coil structures and larger than typical folded proteins, with the exception of two cases (Fig. 4c). In contrast to other works [44, 45], our model has not been directly trained on IDPs; nonetheless, it provides zero-shot predictions of *R*_*g*_ that correlate well with experimental measurements, albeit with overestimation of *R*_*g*_ values for longer sequences (Fig. 4c).

**Fig. 4.**
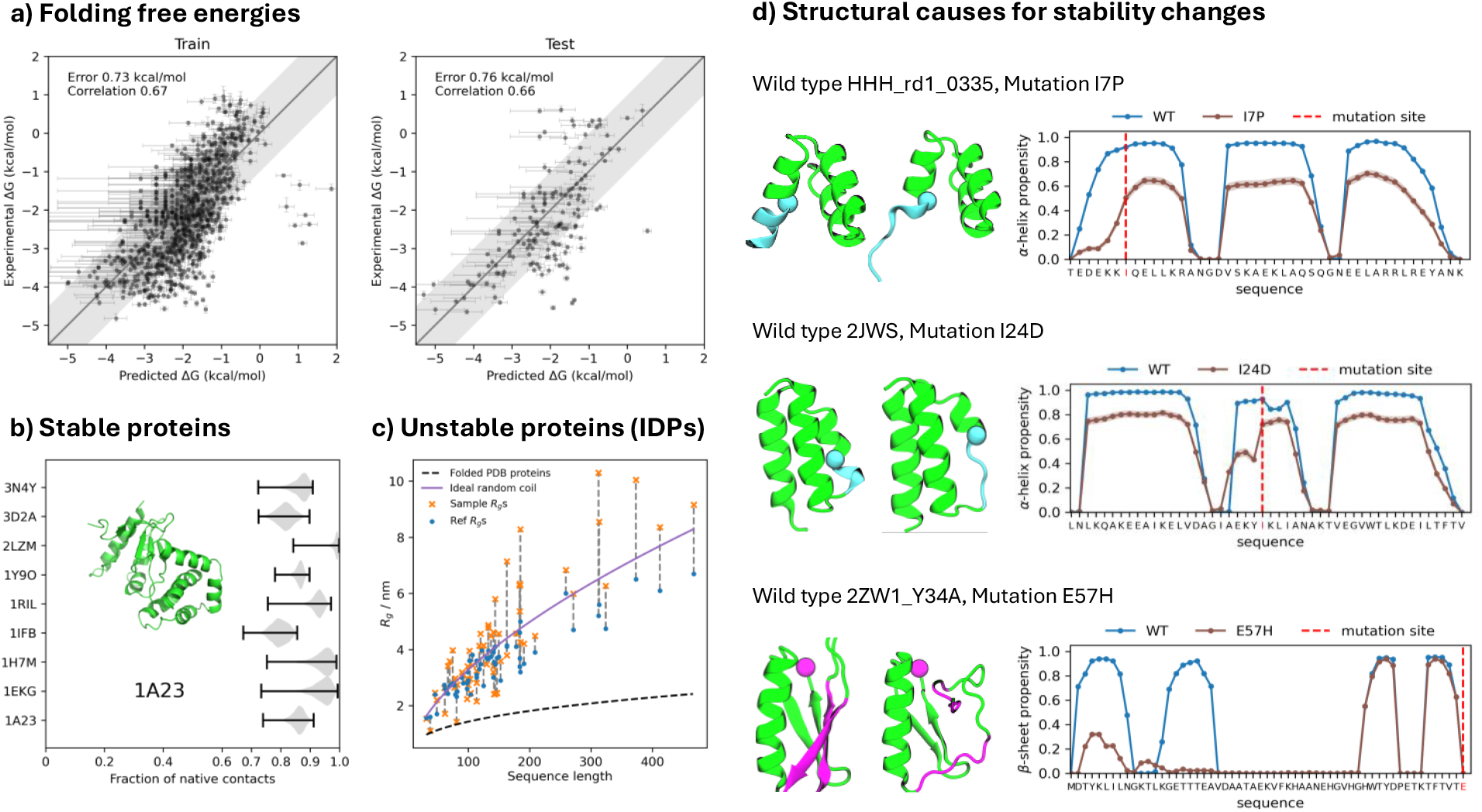
Prediction of experimentally measured protein stabilities. a) Comparison of experimental measurements of folding free energies [21] with model predictions, generated by direct sampling and counting of folded and unfolded states for train and test proteins. b) Validation that very stable proteins that are not included in the MEGAscale experimental dataset are consistently predicted as folded. c) Validated that intrinsically disordered proteins (IDPs) reported in [43] and [45] are predicted as unfolded. Radius of gyration (*R*_*g*_) is compared between model (orange crosses) experimental measurement (blue dots) and Flory scaling [46]. d) Analysis of the effect of three destabilizing mutants on the folded structures as predicted by the model: HHH_rd1_0335 with mutation I7P, 2JWS with mutation I24D, 2ZW1_Y34A with mutation E57H.

In contrast to directly predicting Δ*G* by supervised learning models, we can analyze the structure ensemble generated by our model to reveal insights on mutation-caused stability changes. For illustration, we show mutants of the design protein HHH_rd1_0335 and PDB entries 2JWS and 2ZW1 (Fig. 4d). In HHH_rd1_0335, the mutation I7P leads to a destabilization of the first helix, as indicated by the model’s prediction of a ΔΔ*G* of 1.3 kcal/mol compared to the experimental 2.1 kcal/mol. The analysis shows a decrease in average helicity which particularly affects the helix where the mutation is located. In the case of 2JWS, the mutation I24D in the middle helix results in its partial unfolding, with the model predicting a ΔΔ*G* of 1.5 kcal/mol against an experimental value of 2.9 kcal/mol. This mutation replaces a hydrophobic residue with a negatively charged aspartate, disrupting core stability and leading to a localized structural change. Lastly, for 2ZW1, going from a single mutant Y34A, to a double mutant Y34A & E57H introduces a positive charge in a region surrounded by other positively charged residues, leading to a predicted ΔΔ*G* of 1.9 kcal/mol compared to the experimental 1.2 kcal/mol. This mutation significantly disrupts the N-terminal beta-hairpin, a prediction that aligns with our short MD simulations showing similar destabilization. These analyses highlight BioEmu’s ability to correlate predictions of thermodynamics with structural causes, which is not possible with black-box prediction models.

## 6 Conclusion

We have introduced a generative machine learning system to approximately sample the equilibrium distributions of proteins and through that explore two key aspects of molecular function: protein conformations and their equilibrium probabilities. The system has been demonstrated to sample experimentally known structures of proteins undergoing a variety of conformational changes, to approximate the equilibrium distributions of extensive MD simulations, and to predict experimentally-measured protein stabilities within errors of 1 kcal/mol. The cost of running inference is on the order of one GPU-hour per computational experiment — many orders of magnitude less than running MD simulations even if enhanced sampling methods are invoked, and orders of magnitude cheaper than experiments that can provide detailed structure-function relationships.

BioEmu and MD simulation are complementary: our system was trained on large amounts of MD simulation of soluble proteins, and within this scope, it has shown to be able to approximate MD distributions at a tiny fraction of the MD simulation costs. However, BioEmu cannot be expected to generalize beyond this scope — for example membrane environments and small molecule ligands are neither represented in the model nor in the training, and BioEmu can therefore not be expected to make reliable predictions when membranes or ligands play a key role in the process. For MD, generalizing to such conditions is straightforward, although obtaining results will be limited by the sampling problem.

Our system can be used to generate a guess for the equilibrium distribution, and MD trajectories can be launched from a BioEmu ensemble in order to obtain chemically accurate all-atom structures, refine the distribution, and even compute dynamical properties. We therefore do not expect that emulators such as BioEmu will make MD simulation obsolete; however we do expect that the role of MD simulation will shift from a production tool to a data generation and validation tool, as is already the case for other simulator-emulator pairs such as quantum chemistry and machine-learned forcefields. We have demonstrated that BioEmu can be efficiently fine-tuned on experimental data such as folding free energies. This is an important advantage compared to MD forcefields, which can also be tuned to fit experimental data [47], but the processes that give rise to the experimental observables must be sampled during the training process — a task that is tedious or even unfeasible for observables that involve complex rare events, such as folding free energies.

An important limitation of BioEmu is that it generates distributions entirely empirically, while MD simulation uses potential energy functions which are connected to equilibrium distributions and expectation values by statistical mechanics. If direct access to a potential energy function *u*(*x*) was available that is consistent with the generated distribution by *p*(*x*) ∝ e^*u*(*x*)^, it could be used for reweighting and making rigorous enhanced sampling simulations available through the emulator. Another limitation of the current system is that it only emulates single protein chains at a fixed thermodynamic condition of 300K. A proper emulator for proteins requires conditioning on experimentally and biologically relevant parameters such as temperature and pH, and needs to be able to model multiple interacting molecules, as proteins rarely have a function on their own.

While structure prediction systems for predicting biomolecular complexes already exist [2, 3], a key obstacle to extending the emulator to other scopes different from proteins, as well as further improving it for the current scope, is the lack of training data. While we have shown that the ability to accurately emulate the equilibrium distributions of small proteins increases with more training data, the sampling problem limits MD to generating data for small fragments of biomolecular systems. For learning changes of conformation and binding state of large biomolecular complexes, as well as learning the subtle binding affinity differences between binding partners, and ultimately tackle the quest for reliably predicting protein function, highly scalable experimental techniques that can be incorporated as training data will become key.

## Code availability

The BioEmu model and inference code are available under MIT license at https://github.com/microsoft/bioemu. Code for benchmarking BioEmu or other models for their ability to sample multiple protein conformations, free energy surfaces and folding free energies (Figs. 2, 3b, 4a, and Suppl. Figs. S1, S2, S3, S4), is provided under MIT license at https://github.com/microsoft/bioemu-benchmarks.

## Acknowledgments

We are indebted to the entire Microsoft Research AI for Science team, and thank in particular the following individuals for valuable support and discussions: Chang Liu, Peiran Jin, Tie-Yan Liu, Chris Bishop, Bonnie Kruft, Jonas Köhler, Thijs Vogels, Marwin Segler, Rianne van den Berg, Marco Federici, Stratis Markou, Maik Riechert. Furthermore, we thank colleagues from FU Berlin for valuable discussions and advice, in particular Nicholas E. Charron, Katarina Elez, and Aldo Sayeg Pasos Trejo. Additionally, we thank Gregory R. Bowman, Sukrit Singh, and Anthony M. Trent for their help with Folding@Home data and Nuria Plattner for her support.

## Supplementary Material

### S.1 Data

#### S.1.1 AlphaFoldDB processing

An AlphaFold database (AFDB) snapshot was downloaded in July 2024 and preprocessed for model pretraining (Fig. 1d,e). The aim of this preprocessing is to identify sets of similar sequences with heterogeneous predicted structures, and this is accomplished through a series of steps:

1. We used `mmseqs` [1] to cluster all sequences at 80% sequence identity and 70% coverage, resulting in a set containing more than 93 million clusters.
2. To reduce the sequence clusters to a representative set, we clustered the centroids of the these clusters at 30% sequence identity and discarded all but the 80%-sequence-identity cluster containing the centroid of each 30%-sequence-identity cluster. The result was a set of sequence clusters with 80% sequence similarity within each cluster and at most 30% sequence similarity between the centroids of different clusters.
3. We discarded sequence clusters with fewer than 10 members, leaving roughly 1.4 million sequence clusters.
4. We performed structure-based clustering within each sequence cluster, using `foldseek` [2] (version 9.427df8a) with a sequence identity threshold of 70% at 90% coverage.
5. We discarded everything except the representative member of each structure cluster, leaving a set of sequence clusters which each contain a few structure representatives.
6. We discarded sequence clusters with only one structure representative and those where all the structure representatives were disordered (defined as being composed of more than 50% coil in their secondary structure).
7. To account for structural heterogeneity that was incorrectly flagged due to missing regions in structure representatives, we performed structural alignments in sequence-aligned regions of proteins, and discarded structure representatives with a TM-score greater than 0.9 to another structure representative, as computed by `foldseek`.
8. Similarly to [3], we removed sequence clusters lacking at least one structure with pLDDT greater than 80, and with a pLDDT standard deviation lower than 15 across residues

After running this pipeline, we had ~50k sequence clusters with structural diversity. We utilize the structural diversity to generate augmented data during the pre-training phase (see section S.3.2).

#### S.1.2 Protein Data Bank processing

A snapshot of the PDB was downloaded in Nov. 23rd 2023, including all of the available asymmetric units in the mmCIF format. We use the `pdbecif` Python package (version 1.5) for mmCIF parsing and consider an entry for processing if the overall number of residues in the entry was below 2500 with a resolution below 9.5 Å, when a resolution value was available. All molecular entities per entry were separated according to their type (*i.e*., polymer or non-polymer), discarding those associated with other nucleic acids (*e.g*., RNAs, DNAs). Non-biologically relevant non-polymer entities (*e.g*., solvents, ions) were further filtered out by a list provided in [4]. Non-binding polymer chains were then kept on an entry basis if they contained standard amino acid types and depending on whether other non-binding chains corresponding to the same entity identifier had already been processed. So as to better capture ligand-binding conformational effects, all binding polymer chains with unique binders, as determined by a distance threshold of 6 Å between any binder and protein atom, were kept.

#### S.1.3 Molecular Dynamics simulation data

In the following, we list synthetic all-atom molecular dynamics (MD) data used in this article. An overview of all publicly available and in-house datasets is given in Tab. S1. In-house datasets are described in detail in S.1.5, with our standard MD protocol specified in S.1.4. Public datasets are listed under S.1.6. As our model is for single protein chains and there were several MD simulations of multi-chain systems, we extracted single protein chains from those and treated them independently. As a consequence, the effective cumulative simulation time for such multi-chain simulations is reported as a sum over all chains. For datasets curated from the literature, a reference is provided, whereas details for MD simulations generated specifically for this work can be found in S.1.5.

**Table S1.** Molecular dynamics training datasets used in this work, their associated number of systems, number of individual chains, simulation time, and forcefield used.

| Dataset | Sim. time (ms) | Eff. sim. time (ms) | Force field | # MD sys. | # ind. chains | Ref. |
| --- | --- | --- | --- | --- | --- | --- |
| DESRES-fastfolders | 8.2 | 8.2 | charmm22* | 12 | 12 | [5] |
| FAH-DDR1 | 6.8 | 6.8 | amber ff99sb-ildn | 9 | 9 | [6] |
| FAH-SETD8 | 5.9 | 5.9 | amber ff99sb-ildn | 26 | 26 | [7] |
| FAH-sarscov2 | 4.5 | 4.5 | amber ff14sb | 2 | 2 | [8] |
| FAH-sarscov2-exascale | 56.5 | 81.0 | amber ff03 | 24 | 46 | [9] |
| FUB-MHCII | 8.9 | 26.2 | amber ff99sb | 68 | 201 | [10] |
| FUB-barnase-barstar | 2.0 | 4.0 | amber ff99sb | 2 | 4 | [11] |
| MSR-cath2 | 41.0 | 41.0 | amber ff99sb-ildn | 1040 | 1040 | S.1.5.3 |
| MSR-megasim | 3.8 | 3.8 | amber ff14sb & ff99sb-disp | 271 | 271 | S.1.5.4 |
| MSR-megasim-mutants | 21.5 | 21.5 | amber ff99sb-disp | 21458 | 21458 | S.1.5.4 |
| ONE-cath1 | 5.2 | 5.2 | amber ff99sb-ildn | 50 | 50 | S.1.5.2 |
| ONE-octapeptides | 8.0 | 8.0 | amber ff99sb-ildn | 1100 | 1100 | S.1.5.1 |
| **Total** | 172.2 | 216.0 |  | 24062 | 24219 |  |

#### S.1.4 MD simulation protocol

We internally developed code specifically tailored towards running large MD production campaigns on Azure compute resources. Our code is based on OpenMM [12] as its compute engine, albeit setups are generated using OpenMM or GROMACS [13] as a backend, depending on each case. Unless noted otherwise, we conform to the following protocol for running our MD simulations: We use exlicit solvent and the tip3p water model [14], solvate structures in a cubic box with 1 nm padding and a NaCl buffer of 0.1 M. The solvent is equilibrated with a harmonic constraint force on the solute heavy atoms for 0.1 ns under constant temperature and volume (NVT) followed by 0.9 ns of simulation under constant temperature and pressure (NPT). The constraint force is switched off in multiple steps during another 0.1 ns simulation time. During the equilibration phase, the integration timestep is set to 2 fs. Production runs are conducted in the NPT ensemble, using hydrogen mass repartitioning with hydrogen mass of 4 amu [15], with hydrogen bond constraints, and an integration timestep of 4 fs. The temperature is set to 300 K and the pressure to 1 bar, unless noted otherwise.

Most simulation data described in this work was generated using T4-based (NC4as T4 v3) Azure compute instances.

#### S.1.5 In-house MD datasets

##### S.1.5.1 Octapeptides

The octapeptide dataset consists of 1100 peptides of 8 amino acids length. The selection of systems had been previously described (see Ref. [16] for details about system selection and initial structure seeding procedures). We extend this dataset with longer trajectories in order to obtain a better representation of equilibrium. For each system, 5 new trajectories with 1 *μ*s length were generated using our in-house protocol (S.1.4), using the same force field as in the original dataset (amber ff99SB-ildn [8]), at 300K. The total simulation time amounts to 8 ms.

##### S.1.5.2 CATH1

This data consists of 50 CATH domains as previously described in Ref. [16]. We also extend this dataset, previously consisting of 4 × 0.5 *μ*s trajectories per system, with longer MD trajectories to better represent long-timescale dynamic behavior of different protein domains. Trajectories between 1 and 5 *μ*s length using the amber ff99SB-ildn forcefield [8] were produced, totaling 100 *μ*s per CATH domain. Data production was conducted using an adaptive sampling [17] scheme, where the first trajectory epoch was seeded from a reference PDB structure, while the following 2 epochs were seeded by extracting frames from previous epochs via a minRMSD clustering [18] approach. The total simulation time of the combined dataset (original and in-house) amounts to 5.2 ms.

##### S.1.5.3 CATH2

The CATH2 dataset focuses on sequence coverage rather than overall simulation length. Similar to CATH1 and to the procedure described in [16], systems were selected from the CATH database [19] (version 4.3.0) by filtering out non-contiguous structures or sequences, non-standard amino acids, proteins with disulfide bonds, coil fractions above 50%, and proteins with a relative shape anisotropy ≥ 0.05. Only domains containing between 50 and 200 amino acids were selected, forming a set of ~1100 domains. Out of those, for 1040 we could generate valid MD setups using our in-house protocol (S.1.4). 1 *μ*s trajectories for these domains were generated using the amber99SB-idln [20] forcefield, producing a total of approximately 39 *μ*s per CATH domain, with the exact amount varying due to compute availability reasons. We used the same adaptive sampling strategy as in the CATH1 dataset, and 2 epochs of reseeding. The cumulative simulation time for the whole dataset is 41 ms.

##### S.1.5.4 MEGAsim

The MEGAscale domain simulation dataset (“megasim” in short) is dedicated to including folding-unfolding transitions in the training data. In total, it consists of extended simulations of 271 wildtypes, and 1 *μs* simulations for each of the 22,118 point mutants, including single-residue insertion/deletions. The systems and mutants in our dataset were taken from the megascale measurements of protein domain stability via cDNA display proteolysis [21]. To ensure that every sampled system had a corresponding experimental measurement of the folding free energy Δ*G* during the folding process, we focused on wildtypes and mutants within the curated set (“Dataset2_Dataset3”) of the reference publication. Our final dataset consists of a smaller subset of systems due to finite computational resources and several applied filters, detailed below.

For the wildtype dataset, we tailored the general simulation setup to efficiently sample in both the folded and unfolded states. The seeding structures included both the folded state as well as less structured decoys. Folded structures were obtained from the AF2 predictions available on the Zenodo repository of Ref. [21]. Unfolded (decoy) starting structures were obtained by simulated thermal denaturization in implicit solvent at 400K, followed by several rounds of adaptive sampling in explicit solvent at an elevated temperature. For both the equilibration and production phases, two force fields were used: amber ff14sb [22] and amber ff99SB-disp [23]. In comparison to more traditional force fields like ff14sb, ff99sb-disp is specifically designed to model disordered proteins and does not over-stabilize globular decoy structures [23]. Even though generally reliable, we noticed that a99sb-disp can destabilize the native fold of a protein after extensive simulations. For those cases we relied on ff14sb to generate samples of the folded state.

To optimize compute efficiency, we chose a rhombic dodecahedron simulation box with a 1.5 nm padding for each individual seed. Equilibration was performed with 0.2 ns NVT and 0.6 ns NPT simulations, targeting 295K and 1 bar with a Langevin integrator and a 4 fs time step. Production simulations were performed for 1.5 *μs* per starting structure at 295K in the NVT ensemble and a 4 fs time step. Bond constraints and hydrogen mass were kept identical to Section S.1.4, and we discarded the first 500 ns of each trajectory to only consider the last 1 *μs* in the subsequent analysis. Post processing was carried out with the goal of obtaining a clear separation between folded and unfolded samples as well as minimizing the effect of mixing samples from two force fields. We used the fraction of native contacts (FNC) to define the relative foldedness of each MD frame, and built FNC histograms for all samples from each force field. While, theoretically, for two-state folding-unfolding transitions one can expect the a bimodal distribution, in practice it can be multimodal. However, we observed that the folded and unfolded states have well-defined density peaks in the FNC distribution, and thus performed a kernel density estimation. For each forcefield, we used the FNC with the lowest estimated density as the folding threshold.

For the unfolded state, we picked the samples below the FNC threshold from trajectories simulated with the ff99sb-disp forcefield, whereas for the folded ones we used samples from the same forcefield, i.e., ff99sb-disp by default. However, some cases remained where the samples above the FNC threshold from the ff99sb-disp forcefield were multimodal, that spread over a large range, or that had significant lower FNCs than the samples from ff14sb. For those systems, we selected ff14sb for the folded state. We discarded systems where neither force field resulted in a clear density peak in the FNC distribution above 0.8, or where either the folded or unfolded samples consisted of less than 10% of the entire dataset. In difficult cases, we checked several sample structures as well as the FNC and RMSD time series and made a decision based on visual inspection. After processing, the “MSR-megasim-merge” dataset consisted of 271 wildtype systems, out of which 77 had folded states from amber ff14sb and unfolded states from amber ff99sb-disp, while the rest 194 featured both folded and unfolded samples from amber ff99sb-disp.

Due to the large number of sequences present in the mutant dataset, we could not afford to conduct sampling as thorough as for the wildtypes. Instead, we relied on the presumption that point mutations or single insertion/deletion mainly affect local interactions in the folded state, and only weakly perturb the sample distribution in the unfolded state. Since our model only considers the protein backbone, we re-used the unfolded samples from the wildtypes for all point mutants, and generated MD simulations for all mutants in the folded state. Here we also assumed that the mutant folded state does not deviate completely from the native state of its wildtype, but would only be involved in local rearrangements, such as side-chain repacking. This assumption allowed MD simulations of mutant structures to be seeded from their wildtype folded conformation, as well as the use of the wildtype FNC to probe mutant foldedness. It further means that we can use the FNC defined by the wildtype native contacts to probe the foldedness of the mutants. In practice, we generated the mutant starting structures from their corresponding wildtype reference structure by exchanging the sidechain accordingly and by performing energy minimization. This is followed by a 1 *μ*s simulation for each of the mutants using the amber ff99sb-disp force field. Since we expect the starting structure to be not the exact native structure of the mutant, we anticipated the need for a burn-in period, in which the system can explore a more stable native folded state. To select the length of such period, we split the trajectory into two parts so that the difference of the mean FNCs of each part would be maximized. The part after the burn-in period was then kept for the folded samples, except for situations where the FNC decreased monotonically throughout the simulation.

To validate the combination of samples of each wildtype with its mutants, we considered the impact of including mutant folded samples on the folding threshold for FNC computation. In cases where the previously classified unfolded samples had surpassed the folding threshold, it was no longer possible to define foldedness for the mutant based on its wildtype native contacts. In all other cases, samples were combined, since those coming from wildtype simulations only contributed to the unfolded population. After excluding cases violating the previous two assumptions, we obtained a set containing samples for 21,458 mutants, which we named the “MSR-megasim-mutants-mosaic-disp” dataset.

##### S.1.5.5 Complexin

We have generated a small MD dataset for complexin-2 (Uniprot ID Q6PUV4), which has only been used to qualitatively compare model predictions in Fig. 3c. The simulations were seeded using the AlphaFold2 predicted structure deposited in Uniprot. First, we produced a 5 *μs* trajectory with the Amber ff14sb force field [22] using our standard MD simulation protocol (Sec. S.1.4). Second, we generated dynamics with the Amber ff99sb-disp [23] force field. Here, the setup and equilibration were conducted in GROMACS [13] with initial structures being solvated in a cubic box with 1.2 nm padding and 0.135 molar KCL buffer and the custom ff99sb-disp TIP4P water model. After local energy minimization, the system was equilibrated in 0.1 ns (NVT) and 0.1 ns (NPT). Four production simulations of 1.5 *μs* were conducted in OpenMM using our standard protocol S.1.4.

We evaluated the simulation speed on NVIDIA TitanV to be 200 ns/day for ff14sb and 60 ns/day for ff99sb-disp, the latter being reduced to the more expensive 4-point water model.

#### S.1.6 Public MD datasets

##### DESRES fast-folding proteins

We use the fast folding protein simulations described in Ref. [8] under license. The dataset consists of 12 systems simulated with the charmm22* force field [24], with a cumulative simulation time of 8.2 ms. This dataset has only been used for a separate model whose results are shown in Fig. 3a and S7, but it is not used to obtain any of the other results presented throughout the manuscript.

##### DDR1

Simulations of 9 DDR1 kinases published in Ref. [6](https://osf.io/4r8x2/), with the amber ff99sb-idln [20] forcefield and featuring a cumulative simulation time of 6.8 ms.

##### SETD8

Simulations of methyltransferase SETD8 [7](https://osf.io/2h6p4/), excluding data complexed with small molecules. The dataset consists of 26 systems, has a total simulation length of 5.9 ms and uses the amber ff99sb-idln force field.

##### SARS-CoV-2 exascale

We use the publicly available subset of the data published with Ref. [9], which consists of simulations for 24 systems and uses the amber ff03 force field [25]. The cumulative simulation time is 56.5 ms (when counting by trajectory), or an effective 81 ms (when treating chains independently). The data was downloaded from https://registry.opendata.aws/foldingathome-covid19/ (AWS resource name arn:aws:s3::: fah-public-data-covid19-cryptic-pockets).

##### SARS-CoV-2 non-exascale

Non-glycosylated SARS-CoV-2 RBD data as published by Ref. [8] and downloaded from https://registry.opendata.aws/foldingathome-covid19/ (AWS resource name arn:aws:s3:::fah-public-data-covid19-antibodies). The dataset consists of a single system with 1.9 ms cumulative simulation time and uses the amber ff14sb [22] forcefield.

##### MHC2 peptide simulations

We use the dataset of MHC2 in complex with peptides as published by Ref. [10]. It consists of 68 systems with multiple chains and uses the amber ff99sb forcefield [26]. The cumulative simulation time is 9 ms (when counting by trajectory) or effectively 27 ms (when treating chains independently).

##### Barnase-Barstar

Simulations provided by Ref. [11], consisting of one system with two chains using amber ff99sb [26]. The cumulative simulation time is 2.0 ms (when counting by trajectory) or 4.0 ms effective (when treating chains indepedently). The dataset was downloaded from https://zenodo.org/records/8252423.

#### S.1.7 Experimental thermodynamics data

High-throughput experimental measurements of protein stability at ambient temperature were used to finetune the model in combination with in-house generated datasets. Specifically, we extracted the Δ*G* (“dG ML”) and ΔΔ*G* values (“ddG_ML”) and the corresponding amino acid sequences for wildtypes and mutants within the curated set (“Dataset2_Dataset3”) from the associated data in Ref. [21], which added up to ~776,000 entries.

### S.2 Model architecture

In this section, we describe the architecture of the proposed model and how it is trained. We define BioEmu as a conditional generative model. BioEmu receives as input a protein sequence and generates independent identically-distributed (i.i.d.) samples from the approximated equilibrium distribution over conformations of that protein. The i.i.d. generation of samples can be parallelized across a batch of random seeds, which allows us to approximately explore the equilibrium distribution of protein conformations orders of magnitude faster than standard sequential, correlated molecular dynamics simulations.

#### S.2.1 Protein sequence encoder

The protein sequence *S* is encoded through the protein sequence encoder (Fig. 1b) to compute single and pair representations using a simplified version of AlphaFold2 [27]. Similar to other works [28], we use pre-trained AlphaFold2 [27] sequence representations. We run the AlphaFold2 container with a few changes; we used only the Uniclust30 database [29] (as of August 2018) as reference for multiple sequence alignment construction via `hhblits` [30], completely excluded templates, and removed the AlphaFold2 recycling iterations. During generation, we set the random seed to 0 and use the single and pair embeddings generated by model 3.

As the protein sequence encoder depends on no other variables than the protein sequence *S*, the single and pair embeddings for all proteins used in training and inference are pre-computed once and stored for fast retrieval.

#### S.2.2 Coarse-grained protein structure representation

BioEmu models 3D protein structures with a coarse-grained representation following [31]. Only the backbone heavy atoms of the protein are represented via the *backbone frame* representation introduced in [27]. Similarly to [31], but unlike [27], side-chains and hydrogen atoms are not explicitly modeled by BioEmu.

To convert an all-atom protein conformation to its backbone frame representation for a given residue, we use its C_*α*_ atom coordinate **r** ∈ ℝ^3^ and perform the Gram-Schmidt algorithm on the displacement vectors C_*α*_ → N and C_*α*_ → C. This yields an orthonormal basis which can be represented as a rotation matrix **Q** ∈ SO(3). Repeating this for each residue, we obtain a sequence of position-orientation tuples, 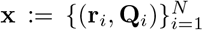, for all *N* protein residues. To recover the Cartesian backbone atom positions from the frame representation, we start with a reference backbone heavy-atom frame per residue type, with idealized atom positions, similarly to AlphaFold2 [27] or OpenFold [32]. For example, for alanine, the idealized frame atom positions are:

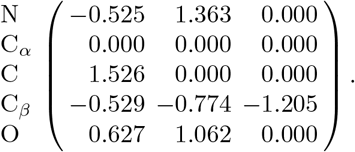

We then apply the rotation matrix **Q**_*n*_ to obtain the rotated frame, and add the position vector **r**_*n*_ to the coordinates of all the atoms in the frame. Note that since the C_*α*_ is at the origin of the idealized frame, it will be at exactly location **r**_*n*_ upon applying this transformation.

#### S.2.3 Diffusion conditional generative model

BioEmu acts as a sequence-conditional generative model: given a protein amino acid sequence, the model parameterizes a distribution of backbone conformational states. Formally, let *S* = (*a*_1_, *a*_2_, …, *a*_*N*_) be a protein sequence with *N* residues *a*_*i*_ ∈ ℛ from the set of 20 standard amino acids. BioEmu is a conditional diffusion model that can be used to sample 3D protein conformations **x** from a conditional distribution

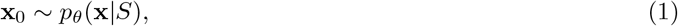

where *θ* are the learnable weights that parameterize the neural network that acts as a score model *s*_*θ*_(**x** *S*). Note that since the dimensionality of **x** depends on the length of the sequence *N*, the dimensionality of the space that BioEmu defines a distribution over depends on the length of *S*. The sampling procedure that characterizes *p*_*θ*_(**x** *S*) is given by simulating the estimated inverse of a *forward diffusion process*, defined by a stochastic differential equation on the space of backbone frame representations **x**:

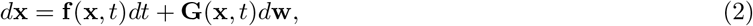

where **w** is a standard Wiener process, and **f** and **G**, drift and diffusion coefficients respectively, are functional hyperparameters. We choose **f** and **G** such that all residues as well as their positions **r** and orientations **Q** are corrupted independently. Specifically, the positions are corrupted with a variance-preserving SDE and a cosine noise schedule as described in [33]. We refer the reader to [34] for further details on diffusing over the space of orientations, SO(3). The orientations are corrupted with a geometric noise schedule so that the marginal distribution of the change in orientation after time *t* is:

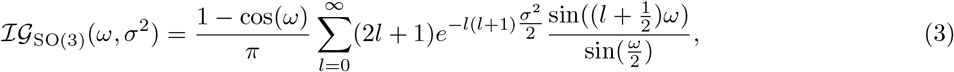

where *ω* is the angle between rotations **Q**_*t*_ and **Q**_0_ computed as:

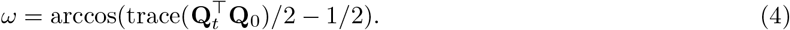

We use *p*(**x**, *t*) to denote the probability distribution of **x** at diffusion time *t* when **x** is corrupted in the above way, with the boundary condition that *p*(**x**, 0) = *p*(**x**) (the target distribution). If the initial positions **r**_0_ are bounded (a reasonable assumption for physical protein structures centered at the origin), then *p*(**x**, 1) is close to a simple *prior distribution* under which positions have a standard isotropic Gaussian distribution and orientations are uniformly distributed.

It has been shown that by training on samples **x**(0) from *p*(**x**) together with corresponding samples from the conditional distribution of **x**(*t*) given **x**(0), a model can approximate the score ∇_**x**_*p*(**x**, *t*); furthermore, if we know the score, we can construct SDEs under which the evolution of the probability density 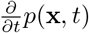 is reversed [35]. Starting by sampling positions **r** and orientations **Q** from the prior and gradually ‘denoising’ by simulating one of these SDEs from *t* = 1 to *t* = 0, we can approximately sample from the target distribution.

Model training details are further described in Sec. S.3. For inference purposes, we smoothen the model weights using an exponential moving average. To sample structures with the trained model, we use the second-order sampler described in [36] with 100 denoising steps, since we found that this resulted in high-quality samples with fewer function evaluations.

#### S.2.4 Score model

The score model (Fig. 1c) takes in single and pair representations of the protein sequence 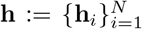 and 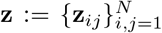, corrupted frames 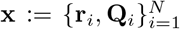, relative sequence positions 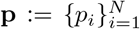, and a diffusion timestep *t*, and predicts the score *s*_*θ*_(**x, h, z**, *t*). It resembles the structure modules of the AlphaFold2 [27] and Distributional Graphormer [28] models, and uses the invariant point attention (IPA) transformer architecture. See Fig. 1c for an overview of the architecture and Algorithm 1 for a detailed description. The translation and rotation scores produced by the score model in Algorithm 1 are defined in the local coordinate frame of each residue, and are invariant under rotation or translation of the entire structure. During denoising, the updates to backbone atom positions are therefore equivariant under rotation and translation of the whole structure.

##### Algorithm 1 Score model *s*_*θ*_(**x, h, z**, *t*)

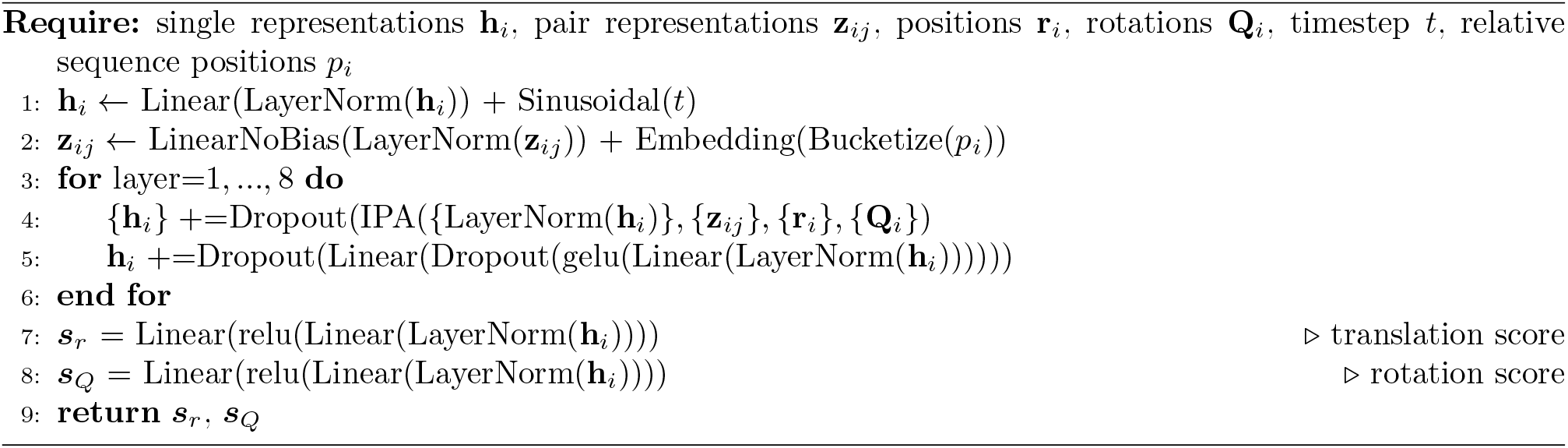

### S.3 Training methodology

We start with a pretrained sequence encoder from AlphaFold2 [27], freeze its weights, and train our own structure module from scratch. We first train on a synthetic dataset derived from AFDB, with high sequence diversity and varied conformations for each sequence (‘AFDB pretraining’ in Table S2, se Sec. S.3.2 for details). The pretrained model can predict diverse conformations for the same protein sequence, but does not quantitatively match the probabilities of different states. We then fine-tune on 95% MD simulation data and folding free energy measurements, mixed with 5% AFDB structures (‘Amber/Δ*G* finetuning’ in Table S2, see Sec. S.3.4 for details), and it results in the main model BioEmu reported in this paper. We also separately finetuned the pretrained model on DESRES fast-folders data (Sec. S.3.3), which was used to produce the results in Figures 3a, S7 but not for any other results.

In all stages, we use the standard denoising score-matching loss as in [31]. To train on experimental thermodynamic data, we add a novel loss term described in S.3.6. Table S2 provides a summary of all training settings. We define a training epoch as the model processing 500K protein structures.

**Table S2.** Training hyperparameters in each stage of training.

| Training stage | AFDB pretraining | Amber/$\Delta G$ finetuning | DESRES finetuning |
| --- | --- | --- | --- |
| Optimizer | Adam | Adam | Adam |
| $\beta_1$ | 0.9 | 0.9 | 0.9 |
| $\beta_2$ | 0.999 | 0.999 | 0.999 |
| $\varepsilon$ | 1e-8 | 1e-8 | 1e-8 |
| Initial LR | 1e-3 | 1e-4 | 1e-4 |
| LR decay factor | None | 0.8 | None |
| LR scheduler patience / epochs | None | 25 | None |
| EMA smoothing factor | 0.995 | 0.999 | 0.995 |
| Max residues per batch per GPU | 2048 | 1024 | 2048 |
| GPU type | A100 | A100 | A100 |
| Number of GPUs | 32 | 4 | 4 |
| Days to train | $\sim 5$ | $\sim 5$ | $\sim 3$ |
| Training epochs | 60 | 100 | 200 |
| Number of seen residues | $\sim 9400\text{M}$ | $\sim 5900\text{M}$ | $\sim 4700\text{M}$ |

#### S.3.1 Data splitting procedure

Having defined a list of test proteins, we removed from our training and validation data any protein whose sequence was similar to any test protein’s sequence. Specifically, we used the `mmseqs2` software [1] (version 15.6f452) and removed proteins if they have 40% or higher sequence similarity with any test protein of at least 20 residues in size, using the highest sensitivity parameter supported by the software (8.0).

#### S.3.2 Pre-training on AFDB

We initially train our model using a dataset derived from AFDB to encourage protein conformational variability (see S.1.1 for details). To draw training examples from this dataset, we randomly select a sequence cluster, and then a structure from within that cluster. While the structure is randomly selected, we always use the sequence associated with the highest pLDDT structure in the cluster as input to the model. This effectively creates a mapping from a sequence to multiple structures (Fig. 1e). In this stage of training, we use the standard denoising diffusion loss [31], defined as a sum over residues. We set the loss to zero in positions where there are insertions or deletions in the sampled structures relative to the representative sequence. The final model checkpoint was chosen based on the performance obtained on our curated OODVal benchmark (see S.4.1). For exact training parameters, refer to table S2.

We compared our pre-training strategy to the more straightforward approach of training the model on the PDB. Additionally, to assess whether the performance of the model was due to an increased diversity in sequence space, we additionally trained on foldseek [2] cluster representatives of AFDB with a pLDDT greater than 90 [37]. This constituted a set of ~250k sequences distinct in both sequence and structure space. We found that models trained on the PDB and the high pLDDT subset of AFDB are significantly worse in its capacity to sample diverse conformations (Fig. S6), indicating that our curated subset of AFDB is an important contribution to facilitate multi-conformational learning.

#### S.3.3 Fine-tuning on CHARMM MD data of fast-folding proteins

For the results shown in Figs. 3a, S7, we fine-tuned our best pre-trained model (S.3.2) on a set of 12 fast-folding proteins [5] simulated with the charmm22* force field featuring sizes ranging from 10 to 80 residues. For evaluation, this set is split into training, validation, and test subsets following a 10:1:1 ratio for each protein (leave-one-out cross-validation). PRB is used as the system used for validation in all splits except in the one where PRB is the test system. In that case, UVF was used instead. Specific training settings are reported in Table S2. All fast-folders results are obtained by evaluated using the model trained in the last available training epoch.

#### S.3.4 Fine-tuning on Amber MD data and experimental folding free energies

Starting from the best model identified during the pre-training stage, we perform fine tuning using the Amber MD datasets listed in Table S1 and described in Sec. S.1.5, plus high-throughput experimental measurements of folding free energies from [21]. In order to retain the performance of the pre-trained models, the fine-tuning data is further augmented with 5 % of randomly-selected AFDB data, with the same settings as in the pre-training stage. The fine-tuned model is trained with hyperparameters shown in Table S2. At this stage, in addition to the standard denoising diffusion loss, for those proteins where this information was available, we also use a novel loss to match experimental folding free energies by backpropagating through the sampling procedure (see S.3.6).

During each training epoch, 500 000 and 50 000 frames are sampled at with a weighted sampler and used for training and validation, respectively. The probability weight of each frame is the product of a *MD dataset weight* and a *normalized frame weight*. The dataset weights, given in Tab. S.3.4, were determined based on accumulated simulation time, sequence diversity and degree of convergence of the different datasets.

**Table S3.**
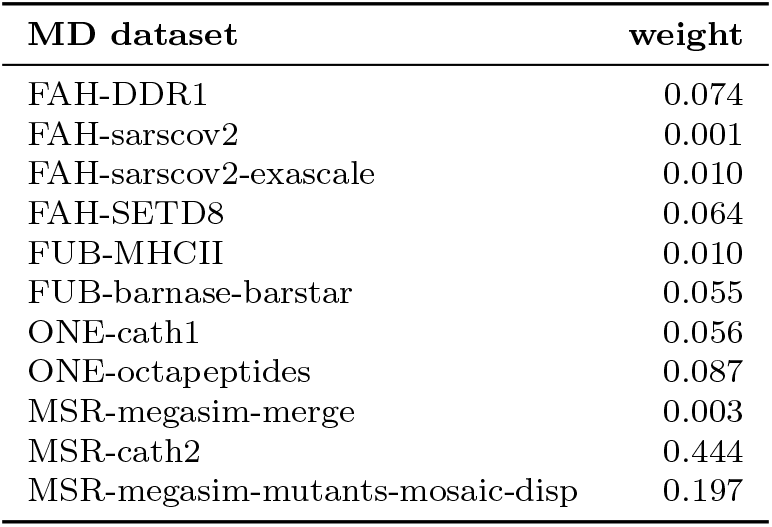
Relative MD dataset weights used for model fine-tuning. These weights define the proportion of samples drawn from a particular dataset.

| MD dataset | weight |
| --- | --- |
| FAH-DDR1 | 0.074 |
| FAH-sarscov2 | 0.001 |
| FAH-sarscov2-exascale | 0.010 |
| FAH-SETD8 | 0.064 |
| FUB-MHCII | 0.010 |
| FUB-barnase-barstar | 0.055 |
| ONE-cath1 | 0.056 |
| ONE-octapeptides | 0.087 |
| MSR-megasim-merge | 0.003 |
| MSR-cath2 | 0.444 |
| MSR-megasim-mutants-mosaic-disp | 0.197 |

Most MD datasets use a frame weight of 1, but specific weights are applied for systems belonging to the following MD datasets:

- Systems in the ONE-octapeptides dataset are reweighted via Markov-State-Models (MSMs) so that each state (a region of configuration space) is sampled with the frequency determined by the MSM equilibrium probability (Sec. S.3.5.1).
- Systems in the MSR-megasim-merge and MSR-megasim-mutants-mosaic-disp datasets are reweighted based on their foldedness so that the training distribution recovers the experimental folding free energies (see S.3.5.3).

In order to deal with systems of varying size, we define batches based on the total number of protein residues in a batch, up 1024 or 2048 depending on the training stage (Table S2). In order to reduce overhead caused by zero-padding, systems of similar size are grouped together when generating batches.

#### S.3.5 Reweighting MD with Markov models and experimental data

To include information about equilibrium properties of the systems in our MD datasets, we have applied different kinds of reweighting to this data, either based on Markov state models (MSMs) or experimental folding free energies.

##### S.3.5.1 MSM reweighting for small peptide datasets

The data distribution generated by MD is often biased towards the seeding structure since simulations are usually run in parallel, often starting from the same or a small number of seed structures. MSMs are a common approach to remedy this problem [38]. In short, the classical approach first projects the 3*N*-dimensional protein system into a low-dimensional representation, discretizes this projection using a clustering algorithm such as *k*-means, and estimates a transition matrix on these discrete states [39]. This approach gives access to the equilibrium probabilities via the eigenvector of that matrix that corresponds to eigenvalue 1. Such eigenvector is then used as a probability distribution to draw samples from the MD simulation accordingly.

This analysis was applied to the ONE-octapeptide dataset, using 2D TICA projections of C_*α*_-C_*α*_ distances and dihedral angles. We used a lag time of 1 ns for both TICA and MSM estimation, and 100 discrete states via *k*-means discretization for the MSM. The obtained equilibrium probabilities are used to draw samples during training.

##### S.3.5.2 Connectivity filtering for post-hoc analyses

It is common practice to perform MSM analyses on sets of states that are reversibly connected. A connected set of states is here referred to as one where each state is reachable from each other state via a sequence of trajectory transitions. Since there can be several connected sets, we choose the connected set with most MD samples in it. As obtaining a connected set from data is numerically more stable than estimating a converged equilibrium distribution, this filter can be applied in situations where a converged MSM estimate could not be obtained.

This analysis was conducted for the ONE-cath1 dataset, based on a linear VAMP projection [56] using a lag-time of 5 ns and residue-residue minimal distances on heavy backbone and C_*β*_ atoms, excluding 1 residue at each terminal and 2 neighboring residues. Subsequent connectivity analysis was conducted by counting transitions between discretized states at a lag time of 500 ns based on the first 5 VAMP dimensions and a *k*-means clustering approach to obtain 200 states. Data outside of the largest connected set was discarded from subsequent analyses, which roughly translated to keeping 90-95% of the data on average. Free energy plots of ONE-cath1 (i.e., CATH domains presented in Fig. 3 and Fig. S8) were based on a secondary TICA projection obtained from trajectories inside the connected set of states.

##### S.3.5.3 Reweighting MD with experimental folding free energies

As detailed in S.1.5.4, we have generated MD simulation data for a subset of the sequences that are represented in the dataset of experimental folding free energies (Δ*G*) provided by [21]. Since the MD simulations are too short to represent a converged sample of folding and unfolding events, the folding free energies estimated from histogramming the raw simulation data do not match their corresponding experimental measurements but mostly correspond to the probability that trajectories were started in folded or unfolded states. To account for this, we reweigh the MD simulation data of each MEGAscale protein system with the corresponding experimental Δ*G*. For each system, first we classify all the MD conformations into folded and unfolded states, and then sample the folded and unfolded structures with different frequencies during training, such that the ratio of folded versus unfolded states seen by the model during training matches the target ratio given by the experimental Δ*G*. Specifically, the folding free energy is related to the probability under the Boltzmann distribution that a protein will be found in a folded state.

The folding free energy Δ*G* can be defined in either of two directions, here we choose the convention to define it as the change in free energy when folding, i.e. Δ*G* = *G*_folded_ − *G*_unfolded_. Then the probability of being in the folded state, *p*_folded_, is related to the folding free energy is Δ*G* by:

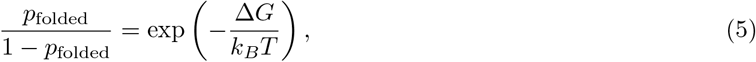

where *T* is the temperature and *k*_*B*_ is the Boltzmann constant. The probability of being in the folded state can be expressed as the expectation value of foldedness *p*_folded_ = *E*_**x**_[*f* (**x**)], where *f* ranges from 0 (unfolded) to 1 (folded). For both our reweighting and model evaluation, we take the form of *f* as:

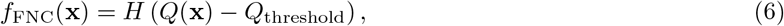

where *H* is the Heaviside step function, *Q*(**x**) the fraction of native contacts, and *Q*_threshold_ a system-dependent threshold. For a given protein structure **x**, the fraction of native contacts (FNC) is defined from pairs of residues that are at least 3 residues apart in the amino acid sequence but which are physically within 10 Å of each other in a reference folded structure. Specifically, we follow notations as in [40]:

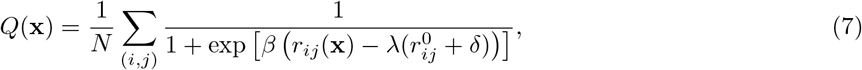

where *r*_*ij*_(**x**) and 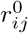 are the contact distances between *i* and *j* in the configuration *x* and the reference conformation (native state). *β* = 5, *λ* = 1.2, and *δ* = 0 are constants, representing the softness of the switching function, the reference distance tolerance and offset, respectively. For each simulated MEGAscale system, we use its PDB structure as the reference conformation. Any given sampled structure can then be classified as folded or unfolded by setting a threshold on the calculated FNC value.

To account for differences in the observed FNC distributions, we set the FNC threshold in a system-dependent but unsupervised manner. Specifically, considering that we initialized multiple MD trajectories separately starting from folded and unfolded states for every protein, and those are not long enough to observe transitions, the distribution of FNC for each system is generally separated into peaks near 1 and 0, representing folded and unfolded states, respectively. In order to obtain a smoother distribution of FNC values for each system, we use a kernel density estimate and then use its minimum within the range of 0.45-0.9.

#### S.3.6 Training on folding free energies via property prediction fine-tuning (PPFT)

Although the reweighting method encourages the model to learn the correct experimental folding free energies with MD simulation data alone, we have empirically found that this convergence is slow, especially for systems where unfolded states are rare (large negative Δ*G*). Even more importantly, experimental observables such as Δ*G* can only be used in a standard diffusion model training approach if folded and unfolded structures are available, e.g. obtained via MD simulation, whose computational costs would limit us to rather few training systems. Here we conducted a large number of MD simulations for 22,389 protein sequences from the MEGAscale dataset, and yet this only corresponds to about 2% of the entire experimental dataset. On the other hand, directly training diffusion models to sample distributions that match a given set of expectation values via generation and backpropagation is computationally prohibitive. The training cost would roughly increase over regular score matching by a factor equal to the number of denoising diffusion steps – in our case that would be a factor 100.

To avoid these limitations and take advantage of high-throughput experiments such as the ones in [21], we have developed a novel and efficient method that trains diffusion models to generated distributions that respect a given set of properties of these distributions, e.g. experimental expectation values. As the method is most likely effective with a pretrained diffusion model, we call it property-prediction fine-tuning (PPFT).

PPFT leverages that many low-dimensional properties of the distribution can be accurately predicted without performing a complete rollout of the diffusion model. Nonetheless, the training principle follows a simple prediction and backpropagation scheme. For a given sequence with an associated experimental Δ*G*, we can roll out the denoising process to generate a clean sample and compute its foldedness. We rewrite Eq. 5 to relate the sample expectation value of foldedness to the folding free energy:

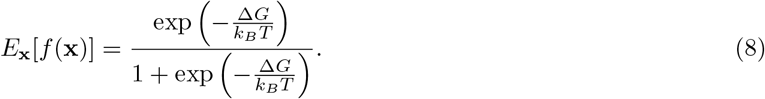

Then, we define the loss term as:

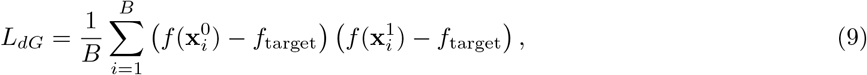

where 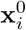 and 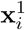 are two i.i.d. samples with the same protein sequence, and *f*_target_ is computed from the right hand side of Eq. 8 using the experimental Δ*G* of the corresponding sequence. The cross term in Eq. 9 is used to minimize the expectation 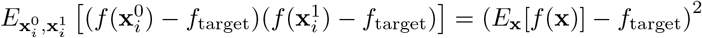. If a standard mean squared error loss were to be used instead, *E*_**x**_ [(*f* (**x**) − *f*_target_)^2^] = (*E*_**x**_[*f* (**x**)] − *f*_target_)^2^ + Var[*f* (**x**)] would be minimized, which contains an additional variance term that would encourage mode collapse.

We notice that the definition of foldedness *f*(**x**) by Eq. 6 is non-differentiable due to the Heaviside step function, and the system-dependent threshold adds additional complication. In PPFT, we instead use foldedness with the following definition:

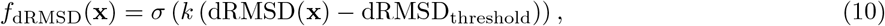

where dRMSD = RMSD(*D*(**x**), *D*(**x**_0_)) refers to the root mean square distance between distance matrices *D*(**x**) of the current structure **x** with respect to a reference structure **x**_0_ (which is neglected in the notation for simplicity). We choose *k* = −24, dRMSD_threshold_ = 0.4 for all protein systems. To enable backpropagation, we use a sigmoid function *σ*, which is differentiable and approaches a Heaviside step function when the slope *k* is sufficiently large.

In the model finetuning stage, for those systems with both simulation and experimental Δ*G* data, we combined *L*_*dG*_ with the usual score matching loss, i.e.:

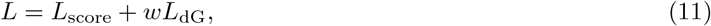

with a weight *w* = 2. Even though the simulation data was reweighted using the experimental Δ*G* based on the method described in Sec. S.3.4, we find that the inclusion of the *L*_dG_ loss significantly sped up Δ*G* model convergence.

As described above, a key requirement for PPFT to be computationally efficient is to avoid executing full diffusion model denoising with hundreds of denoising steps. To mitigate this cost, and considering that folding/unfolding are changes easily recognizable at a coarsed-grained level at earlier denoising levels, we considered reducing the number of integration timesteps, which sacrifices sample quality, but still predicts the foldedness accurately. In practice, we find that 35 timesteps are sufficient when used alongside the Heun sampler. To further reduce cost, we denoised to a specified intermediate noise level *t* to then perform clean sample extrapolation 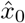 using the reparameterization trick [41], with

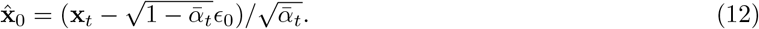

The foldedness is then predicted from 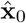 after denoising 8 out of 35 timesteps. We remark that while the coarse-grained nature of foldedness enables us to greatly reduce the number of rollout steps and model evaluations, this may not be applicable for every properties of interest. In such scenarios the adjoint method may be needed for computationally-affordable and numerically-stable training [42].

As a final measure towards increasing efficiency we leverage partial backpropagation, which has shown to effectively reduce computational costs in image-related tasks [43, 44]. Here we apply backpropagation only through the final extrapolation and first 3 denoising steps, i.e., in other denoising steps the score function is detached from the computational graph. We also perform gradient accumulation to effectively increase batch size such that the property prediction loss *L*_*dG*_, as an average over the batch, is closer to the actual expectation value.

To summarize, a combination of techniques contribute to the performance of PPFT:

1. Definition of a differentiable target function,
2. Cross-target matching loss term,
3. Joint training with regular score matching,
4. Gradient accumulation,
5. Use of a higher order sampler to reduce integration timesteps
6. Extrapolation, and
7. Partial backpropagation.

We find that 2-4) are particularly helpful for reducing the mode collapse problem that results from overoptimization of the property prediction loss function, while 5-7) help to greatly reduce the rollout and backprop steps, such that direct backpropagation is feasible with current compute requirements.

### S.4 Multi-conformation benchmarking

We begin by describing which sets, as well as their rationale for inclusion and curation considerations in S.4.1. Additional details about how we constructed an uncontaminated benchmark for evaluation is provided in S.4.2. In S.4.3 we provide details on the summary metrics we use in order to evaluate multiconformational capabilities of our and other competing models. Finally, in S.4.4 we provide insights into the baselines we considered to compare against our models as well as the parameters that were chosen for them.

#### S.4.1 Benchmark sets

In order to evaluate the multiconformation sampling capabilities of the pre-trained and fine-tuned models, we manually curated several sets of examples that are interesting from a structural biology point of view. For some of the benchmarks, this curation also optionally includes manual labeling of residues where a specific conformational change of interest happens. Full lists containing PDB and chain identifiers (`label_asym_id`), as well as residue labels for both alignment and metric computation regions, where appropriate, are provided in the Supplementary Data of this manuscript. Details on each individual benchmark are provided below:

- **OOD60**: A collection of 19 examples collected from the PDB after the AlphaFold 2 monomer model cutoff date (Apr. 30th 2018). A 60% sequence similarity cutoff is used to remove anything from this benchmark that is similar to any chain in the PDB prior to the specified cutoff date. This benchmark represents an unbiased evaluation of multiconformation sampling, and contains several representative examples of the type of conformational changes present in other benchmarks. Metric-wise, we use RMSD as defined on either a local region, or the entirety of the protein, depending on each case.
- **Domain motion**: 22 examples representing large-scale hinge motions. Only global RMSD is used for evaluation in this benchmark.
- **Cryptic pocket**: 34 example pairs featuring a conformational change characterized by the formation of a binding site that is induced in a *holo* (bound) structure, but not on its *apo* (unbound) version. Many of these examples were further curated from the CryptoSite benchmark [45] or other related works [46]. The binding site and other parts involved in the conformational changes were manually-defined, and local RMSD was used as a metric.
- **Local unfolding**: A set of 21 examples, where a certain chain segment of at least 8 residues undergoes an unfolding transition, including some examples from the benchmark proposed in [47]. For this benchmark we defined the segment of the protein that can unfold or detach and measure the fraction of native contacts between this segment and the entire protein to track whether a sample was folded or unfolded.
- **OODVal**: A manually-curated set of 11 examples picked after the AlphaFold 2 monomer model cutoff date but that is disjoint from the OOD60 set detailed above, which we use for pre-trained model selection purposes. Only global RMSD is used as a metric in this benchmark.

#### S.4.2 Curation of the OOD family of benchmarks

As mentioned in S.4.1, we selected pairs of references after the AlphaFold 2 monomer model cutoff date to account for potential dataset contamination at evaluation time. We first extracted and separated all chains for all PDB entries after the mentioned cutoff date. Each individual protein entity inside each entry is then associated with a unique Uniprot segment via SIFTS annotations [48]. A sequence clustering procedure using `mmseqs2` is then applied on all Uniprot segments, using a minimum sequence identity threshold of 0.99. Within each sequence cluster, we perform a structure clustering procedure on the associated PDB entities, similar to the one reported in [49]. This included TM-score as the main comparison metric per sequence cluster followed by an agglomerative clustering procedure as implemented in `scikit-learn`, with a maximum allowed TM-score between clusters of 0.7. Benchmark pairs were selected amongst arbitrarily selected cluster representatives, as long as both members had a minimum resolved sequence length of 50 residues, a maximum fraction of coil residues of 0.4, a minimum shared sequence identity between resolved sequences of 0.8, and a maximum resolved sequence length difference of 50 residues.

For the OOD60 benchmark, care was taken that the remaining pairs were at most 60% sequence-similar to the AFDB training set. OODVal was selected as the set difference between the whole OOD and OOD60 sets. Both sets underwent significant manual curation to ensure unphysical or unrealistic examples were excluded. Some of the criteria applied for curation included checking whether an intra-domain conformational transition was present, whether that occurred in a region that is resolved in both references, or filtering for chains that formed a single long helix, as we deemed their stability outside a complex unlikely.

#### S.4.3 Measuring multiple conformations

For most benchmarks we used RMSD on the backbone atoms as our main metric. For the local unfolding benchmark, however, we used a C_*α*_-only version of a contact map between the unfolding region and the entire protein because the unfolded state has no single reference. When sampling for most multi-conformation benchmarks, we always used the experimental sequence (`_entity_poly.pdbx_seq_one_letter_code_can` in the mmCIF dictionary entry). In cases where the experimental sequence differed between two deposited references, both were sampled in equal proportion. For the cryptic pocket benchmark, however, only the experimental sequence of the *apo* conformation was sampled, as it is the more biologically challenging case. Global pairwise sequence alignments were used to compute metrics as needed when the sampled sequences differed from the experimentally-resolved reference sequences in the mmCIF files. For this, we mostly used the default parameters of BioPython’s `PairwiseAligner`, apart from manually setting an open gap penalty of 0.5.

We computed two key summary statistics in order to evaluate the multiconformation capabilities of our model, as well as to compare it against other approaches:

- **Coverage**: measures the fraction of sampled reference conformations, according to a chosen metric, and as a function of different metric thresholds. We consider a conformation as covered if at least 0.1% of samples are within a specific threshold the corresponding reference structure.
- *k***-recall**: defined as the average of a metric for the closest 0.1% samples per reference.

Before metrics were computed, a filtering procedure was undertaken to discard unphysical samples, both in terms of chain breaks and clashes. Specifically, we looked at C*α*-C*α* and C-N distances between sequence-adjacent residues and ensured that these do not surpassed 4.5Å and 2.0Å thresholds, respectively. Additionally, distances were computed between any two backbone atoms of different residues and we ensured that samples did not contain any such distances below a threshold of 1.0Å. Exhaustive evaluation metrics and summary statistics for these benchmarks is provided in Figs S2-S4, and Table S4.

#### S.4.4 Baseline methods

We chose AFCluster [50] and AlphaFlow [49] as baseline methods for multiple conformation generation. AFCluster is a method that relies on MSA-subsampling techniques in conjunction with AlphaFold2 to generate distinct samples, whereas AlphaFlow, similar to this work, is a deep-learning-based generative model. For both of these methods, MSAs were generated via using ColabFold [51] using the default parameters. In the case of AlphaFlow, the same number of samples were drawn as for our model, whereas for AFCluster, the number of samples was limited to the number of clustered MSAs generated by the method. AlphaFlow runs included the recommended `--self cond --resample` flags when evaluated. Comparisons of our trained models against these baselines on the proposed benchmarks is provided on Fig. S5.

### S.5 Protein stability benchmarks

#### S.5.1 System selection

We selected proteins from ProThermDB [52] such that their experimental Δ*G* of unfolding ≥ 6 kcal/mol and their asymmetric unit contains a single chain. We removed proteins with following conditions, arriving at 26 proteins for the benchmark: several proteins, including two annotated as membrane proteins, one whose sequence was undetermined, and one that was a nucleic acid-protein complex. The initial selection comprised of 140 systems. Additionally, we curate a smaller subset consisting of 26 proteins after excluding systems with one of the following conditions:

- proteins annotated as membrane protein
- proteins with a ligand reported under `_refine_hist.pdbx_number_atoms_ligand` (e.g., 1C52)
- proteins with disulfide bonds as reported under `_struct_conn.conn_type_id` (e.g., 1LVE)
- oligomeric proteins (e.g., 1ROP, supposedly only stable as a dimer) or proteins in protein-RNA complexes
- proteins with ligands not reported in `_refine_hist.pdbx_number_atoms_ligand` (e.g., 2LCP)
- proteins with repeated entries due to differing capitalization

We also use the intrinsically disordered proteins (IDPs) of the CALVADOS test set of IDRome [53, 54] to benchmark stability, which features 65 IDPs. Sequence similarity search indicated that there was only one protein with a similarity above 40% with respect to the training set of BioEmu.

#### S.5.2 Evaluating free energy predictions

We compute the ΔΔ*G* of a mutant using the difference between its Δ*G* with respect to its wild type (ΔΔ*G*_mut_ = Δ*G*_mut_ − Δ*G*_wt_). In order to estimate confidence intervals in the predictions, we used the the Clopper-Pearson method.

### S.6 Energy landscape MAE

Assessing the mean absolute error on protein conformations is a non-trivial task for two reasons. a) Conformational landscapes and corresponding free energy surfaces cannot be directly assessed in 3*N*-dimensional space directly, but require a projection space in which the density, and thus the free energy, are computed. b) Free energy landscapes are often very rough and transitional regions have extremely low probabilities compared to metastable states. The error that a model makes in predicting the relative probabilities of metastable states with respect to each other vs. the probabilities of transition regions can be regarded as two different classes of error. In this paper, our goal was to sample metastable states such as folded or unfolded in the correct ratio, and therefore we focused on the first error. We have chosen the following approach to quantifying the mean absolute error (MAE) of protein free energy landscapes over metastable protein states, which are often referred to as macrostates: First, we parameterized a linear TICA projection (time-lagged independent component analysis [55]). Since TICA, like all dimensionality reduction techniques, is fundamentally limited by availability of data, we have limited this analysis to a subset of test systems with sufficient MD data and to the trajectories within the connected sets described in Sec. S.3.5.2. Second, we have chosen macrostates in the 2-dimensional TICA space, and clustered using Hidden Markov models (HMMs) [56], a commonly-used approach in the analysis of MD simulations. HMMs were estimated at comparably short lagtimes of 1ns and with 3 hidden states as a numerically stable choice.

The macrostate MAE (mMAE) was computed by assessing the relative free energies *G*_*i*_ within each macrostate *i* by sample counting:

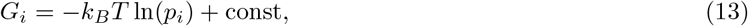

with *k*_*B*_ the Boltzmann constant, *T* the temperature, and *p*_*i*_ the normalized histogram count for macrostate *i*. As not all macrostates were sampled by our model for the systems considered, a prior count of 1 was assigned to each macrostate. For a model with 10k samples that corresponds to clamping *p*_*i*_ = max(*p*_*i*_, 10^−4^), which can be regarded as the model resolution boundary. Relative free energies from ground truth MD distribution and model samples are offset such that min_*i*_ *G*_*i*_ = 0. The overall mMAE between model prediction (ML) and ground truth (GT) was then computed as:

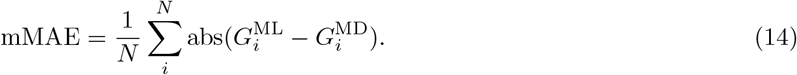

To evaluate BioEmu’s performance on MD-generated free energy landscapes, we have applied our mMAE metric to a random test set from the ONE-cath1 dataset (S.1.5.2). All of these systems have *>* 100*μs* MD data.

### S.7 Supplementary Tables and Figures

**Table S4.** Success rates of multi-conformation benchmark split by whether the reference was present in the AlphaFold2 monomer model training set, or whether it is a new reference not explicitly leaked via embedding poisoning. n_success_ is the expectation value of the number of successful predictions computed via bootstrap, and success is defined for each type of conformational change as described in the main text - domain motion: RMSD *≤* 3Å, local unfolding: fraction of native contacts *≤* 0.3 and *≥* 0.7, cryptic pockets RMSD *≤* 1.5Å.

| **Benchmark** | **Pre-AF2 cutoff** |  | **Post-AF2 cutoff** |  |
| --- | --- | --- | --- | --- |
| | $n_{\text{success}}$ | $N$ | $n_{\text{success}}$ | $N$ |
| Domain motion | 23.89 | 26 | 13.64 | 18 |
| Local folded | 13.16 | 18 | 1.00 | 2 |
| Local unfolded | 9.00 | 11 | 6.30 | 9 |
| Cryptic apo | 17.00 | 34 | 0.00 | 0 |
| Cryptic holo. | 26.96 | 32 | 2.00 | 2 |
| **Total** | 90.01 | 121 | 22.94 | 31 |
| **Success rate** | **0.744** |  | **0.740** |  |

**Fig. S1.**
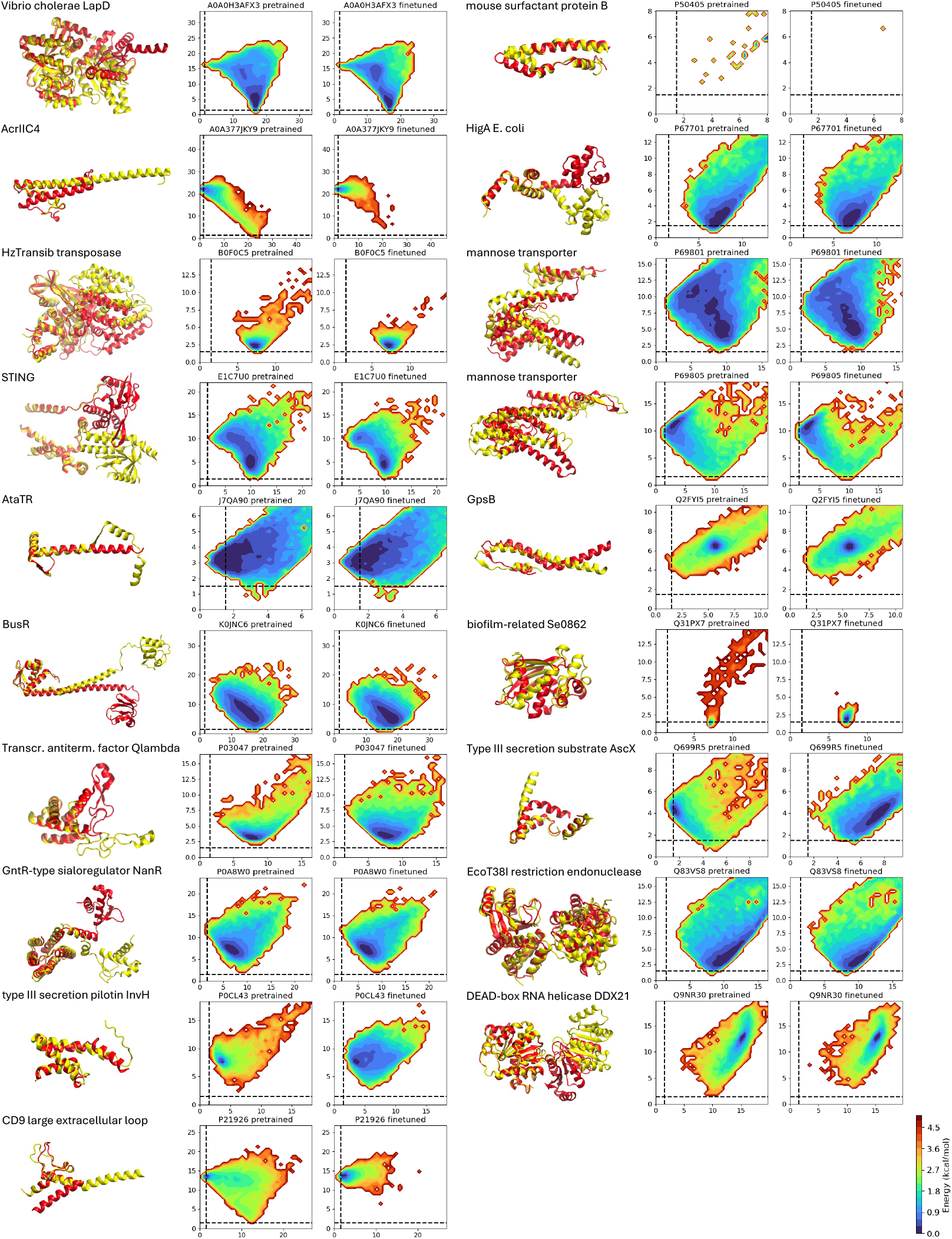
Multi-conformation benchmark: OOD60, i.e., conformational changes of proteins with sequence similarity ≤ 60% compared with the AlphaFold2 training set. For each case, the two reference PDB structures are shown in red and yellow. Energy landscapes show the empirical free energy sampled by the pre-trained and fine-tuned model, respectively, as a function of global C_*α*_ root mean square deviation (RMSD) to each reference. We consider RMSDs below 3Å (dashed lines) as a successful match to the reference structures.

**Fig. S2.**
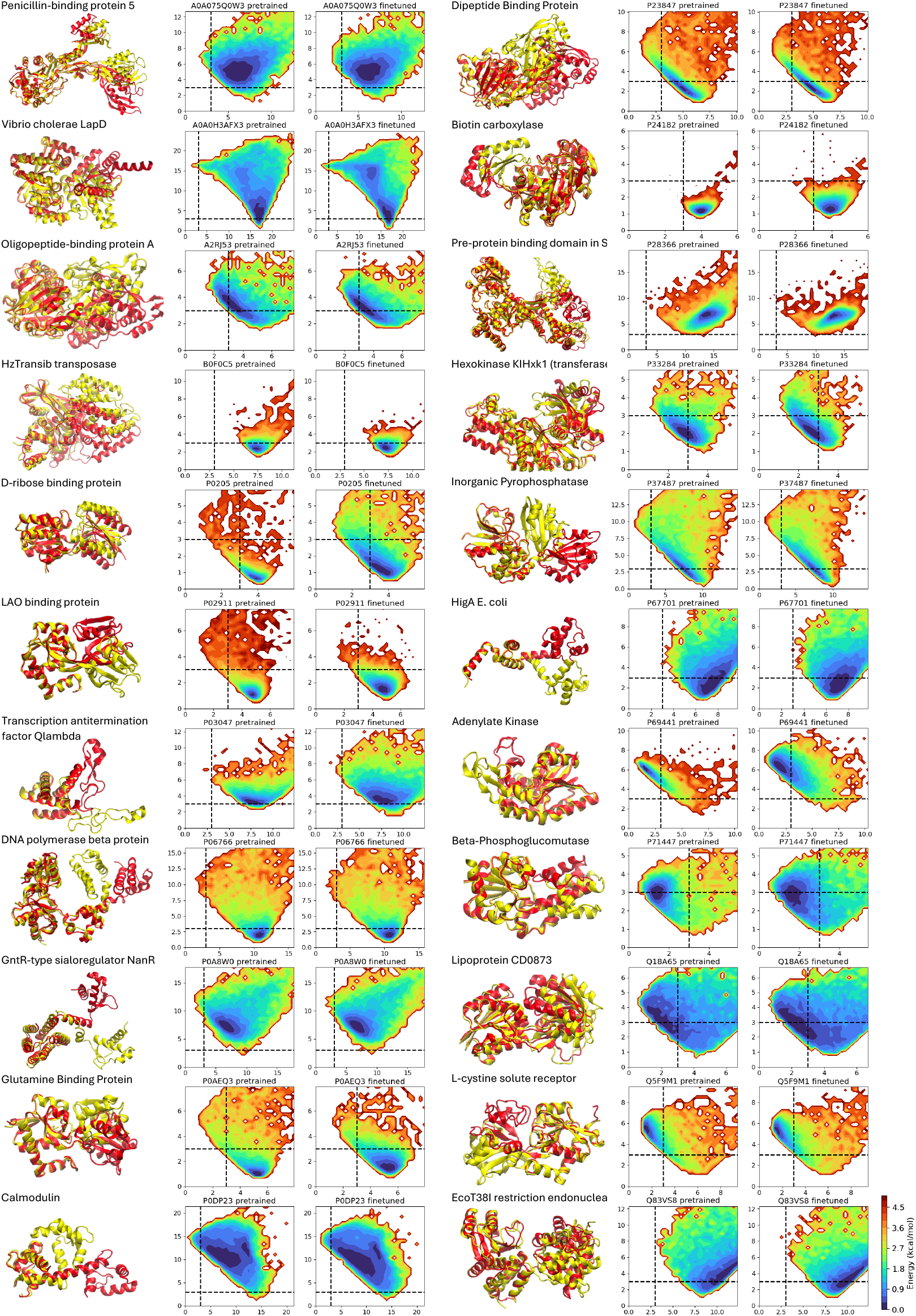
Multi-conformation benchmark: Domain motions. For each case, the two reference PDB structures are shown in red and yellow. Energy landscapes show the empirical free energy sampled by the pre-trained and fine-tuned model, respectively, as a function of global C_*α*_ root mean square deviation (RMSD) to each reference. We consider RMSDs below 3Å (dashed lines) as a successful match to the corresponding reference structure.

**Fig. S3.**
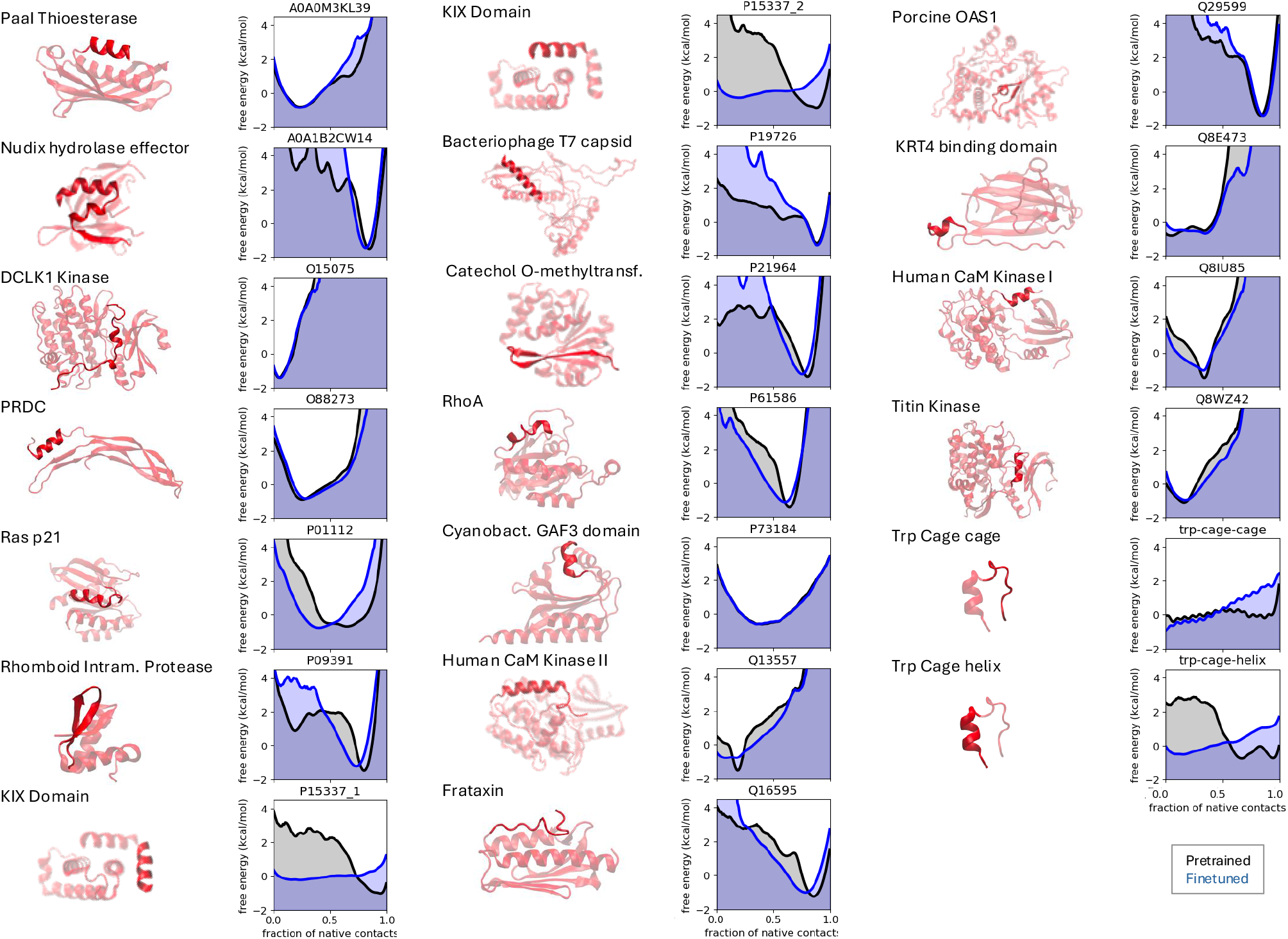
Multi-conformation benchmark: Local unfolding. For each case, the PDB structure used as folded state reference is shown in red, with the part can unfold highlighted. Energy landscapes show the empirical free energy sampled by the pre-trained (black) and fine-tuned (blue) model, respectively, as a function of the fraction of native contacts (FNC) between the region that unfolds with the entire protein. We consider samples with FNCs *>* 0.7 and *<* 0.3 as folded and unfolded states, respectively.

**Fig. S4.**
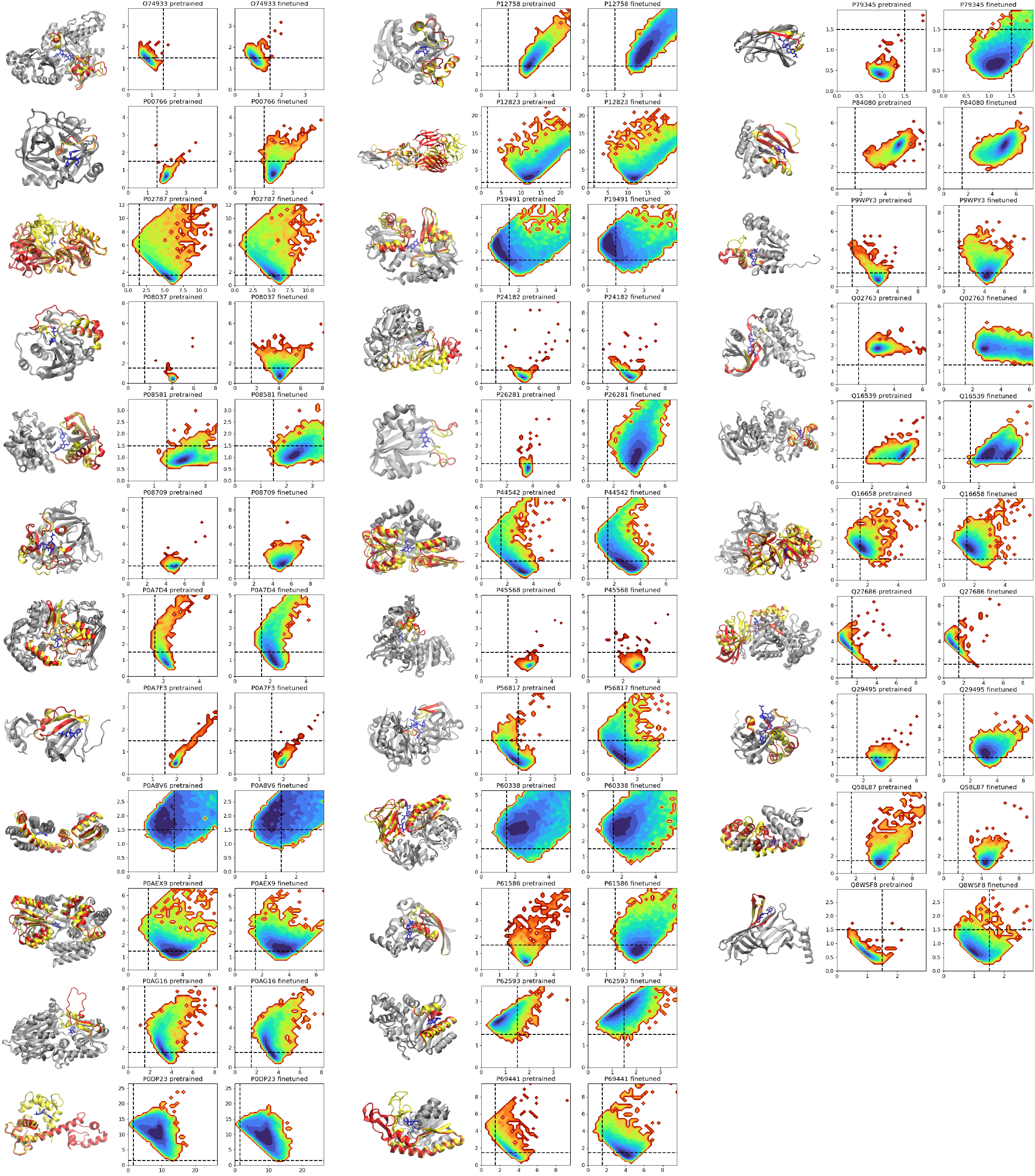
Multi-conformation benchmark: Cryptic pockets. For each case, the two reference PDB structures are shown in red (apo) and yellow (yellow). The holo state residues in contact with the ligand are colored black. Energy landscapes show the empirical free energy sampled by the pre-trained and fine-tuned model, respectively, as a function of a local C_*α*_ root mean square deviation (RMSD) of the region undergoing conformational change, to each reference. We consider RMSDs below 1.5Å (dashed lines) as a successful match to the reference structures.

**Fig. S5.**
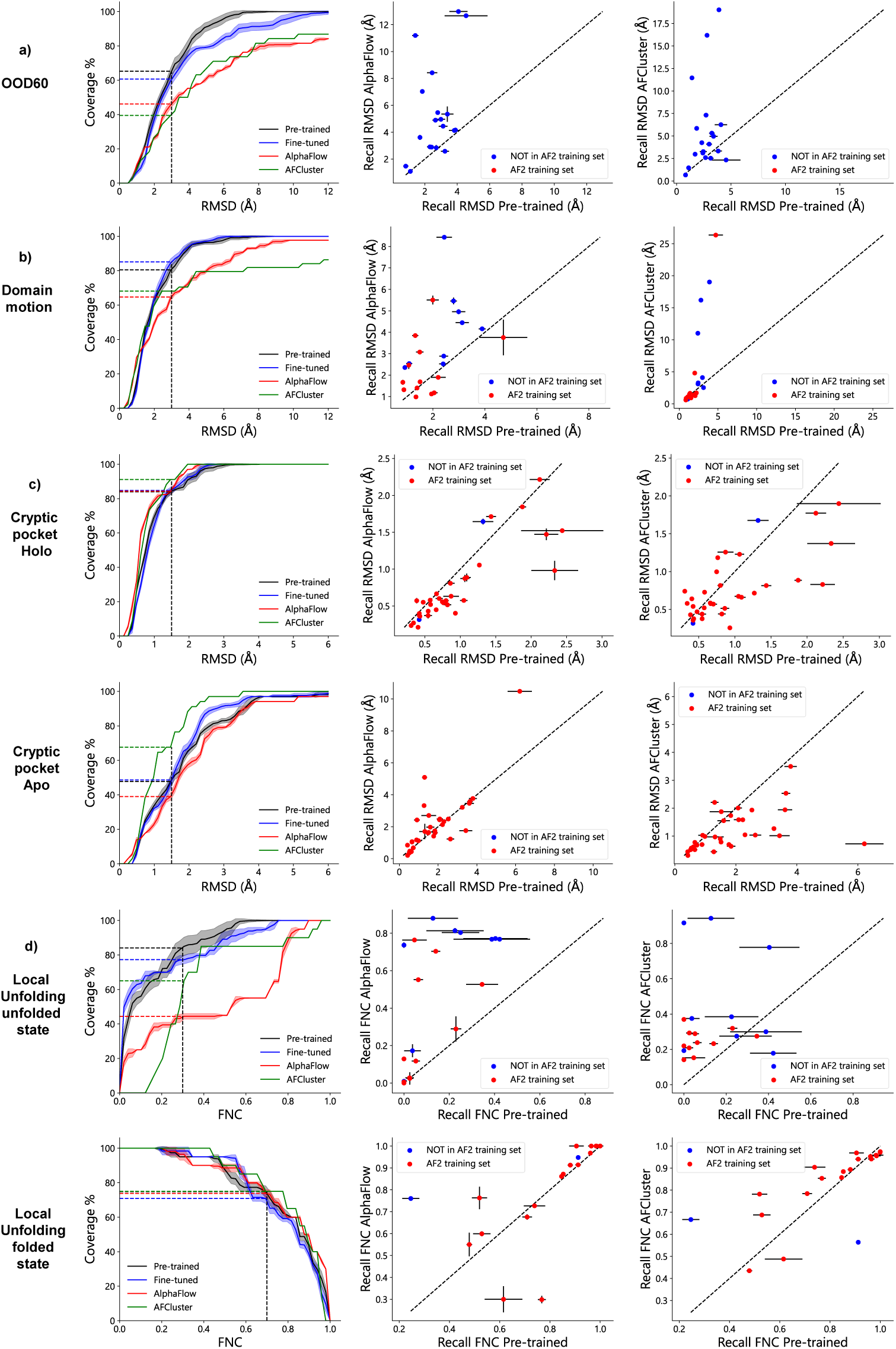
Multi-conformation benchmarking against AlphaFlow and AFCluster for all cases studied in Fig. 2 plus the OOD60 benchmark. Left column: Percentage of reference states covered at a given distance from the reference (higher is better). Middle and right columns: comparison of benchmark-specific metrics for individual benchmark entries between our method (horizontal axis) and AlphaFlow or AFCluster (vertical axis). Note that all comparisons apart from those in the OOD60 benchmark test how well different models fit the data but it is not fair in terms of generalization. Apart from potential Evoformer embedding leakage, for our method, all cases are in the test set, whereas for other methods cases before the AF cutoff date were present in the training set. Our method clearly outperforms AlphaFlow and AFCluster in OOD60, Domain motion and local unfolding, and is particularly strong when generalization is required (blue bullets). AFCluster, on the other hand, significantly outperforms our method on the cryptic pocket benchmark, especially for sampling apo states.

**Fig. S6.**
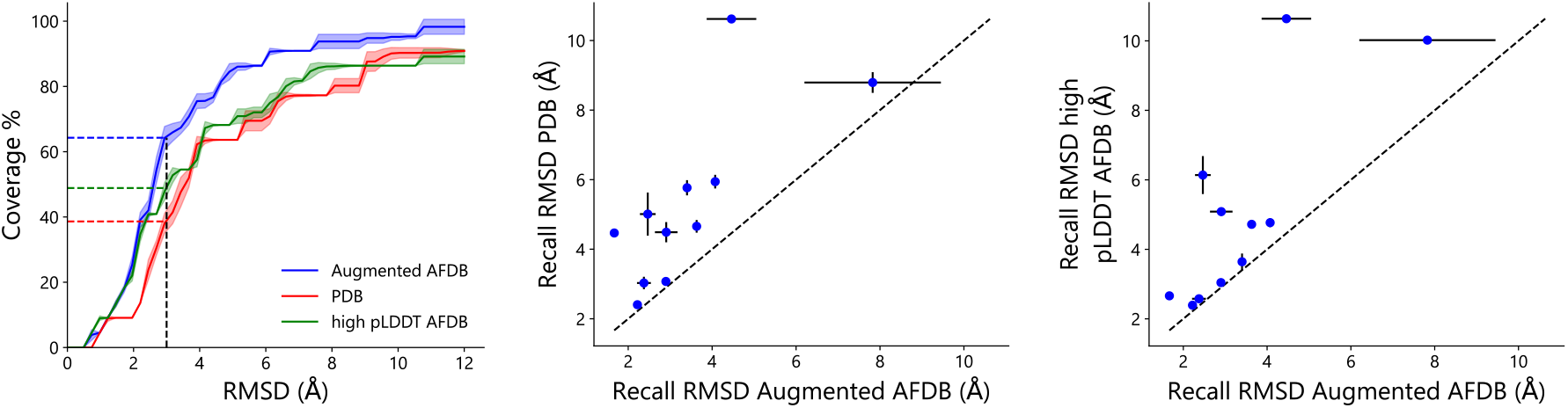
Multi-conformation benchmarking using different pretraining datasets on the OODVal benchmark. Left column: Percentage of reference states covered at a given distance from the reference (higher is better). Middle and right columns: comparison of RMSD values for individual benchmark entries using Augmented AFDB pretraining (horizontal axis) or PDB/high pLDDT AFDB training (vertical axis).

**Fig. S7.**
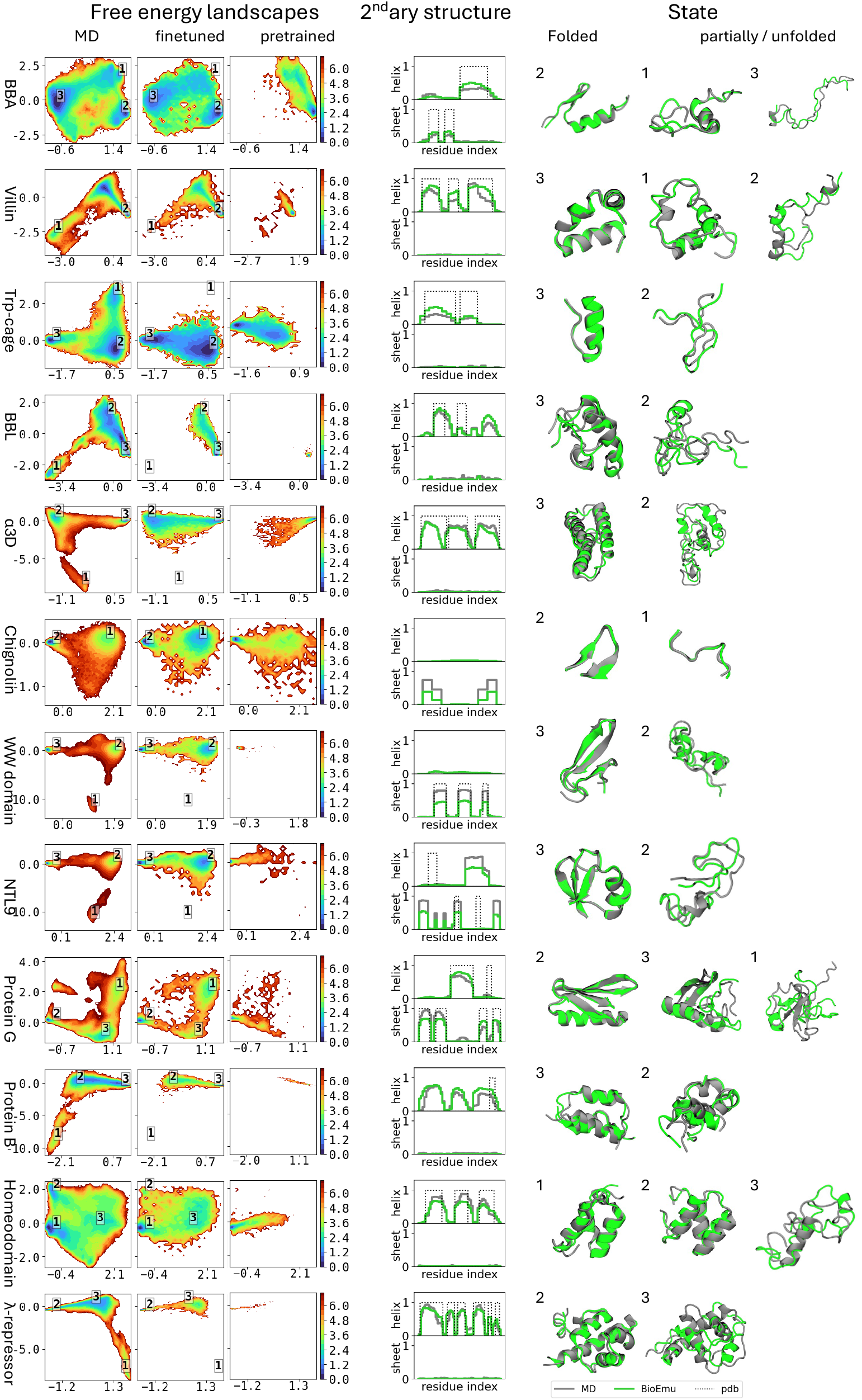
Free energy surfaces for the DESRES fast folding proteins extracted from MD simulations (left), fine-tuned models (center), and pre-trained model (right). Representative structures from MD (grey) and their closest counterparts from the fine-tuned model (green) are shown where BioEmu predicts a state.

**Fig. S8.**
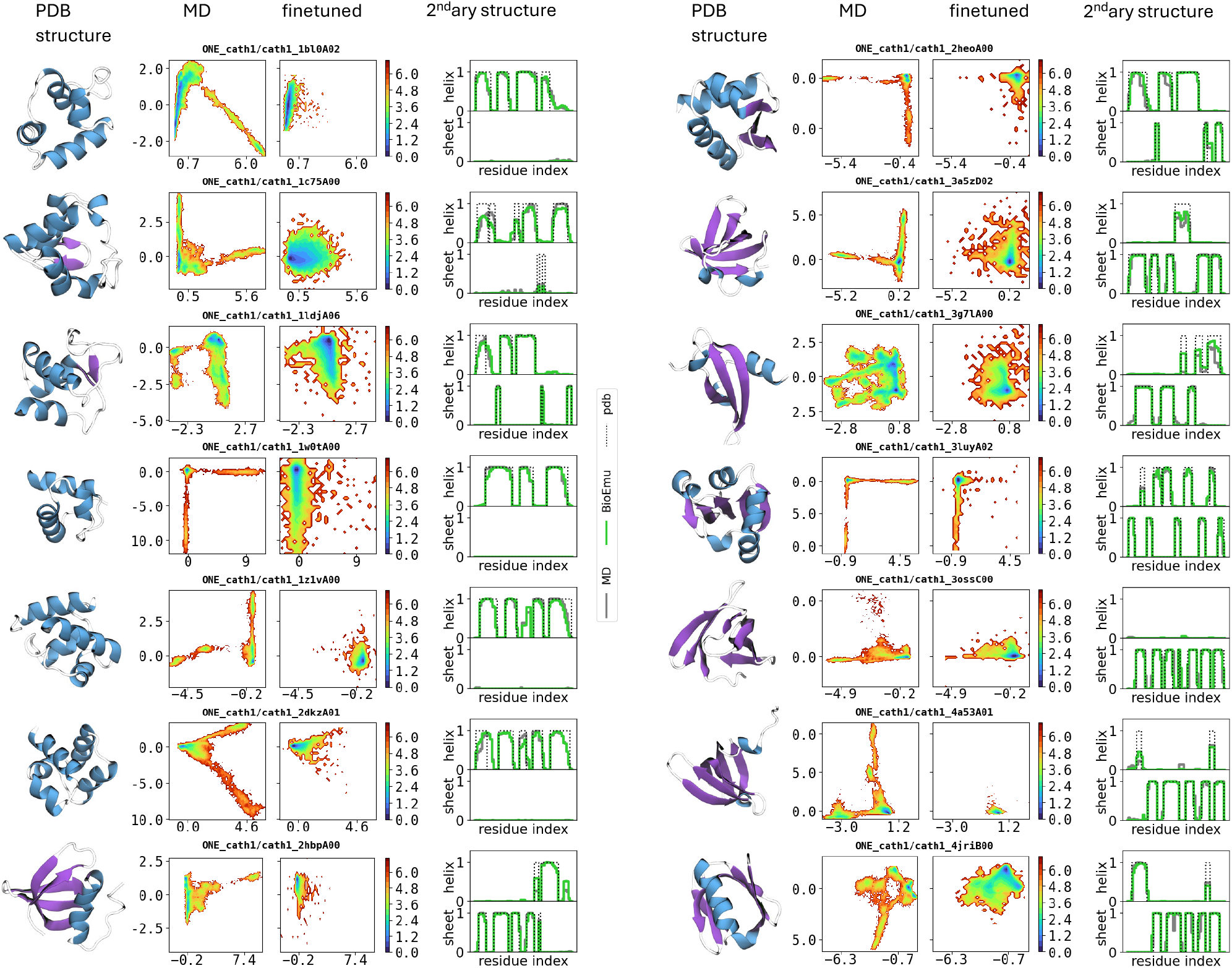
Free energy surfaces for CATH domains of with *>* 100*μ*s simulation time and their secondary structure assignments. Energy surfaces are extracted from MD simulations (left column) and fine-tuned model (center column), respectively, secondary structure assignments (right column) shown as averages.

## References

[1] Jumper, J. et al. Highly accurate protein structure prediction with AlphaFold. Nature, 596(7873):583–589, 2021.

[2] Abramson, J. et al. Accurate structure prediction of biomolecular interactions with AlphaFold 3. Nature, 630:493–500, 2024.

[3] Krishna, R. et al. Generalized biomolecular modeling and design with RoseTTAFold All-Atom. Science, 384:eadl2528, 2024.

[4] Baek, M. et al. Accurate prediction of protein structures and interactions using a three-track neural network. Science, 373:871–876, 2021.

[5] Berman, H.M. et al. The Protein Data Bank. Nucl. Acids Res., 28:235–242, 2000.

[6] Ritort, F. Single-molecule experiments in biological physics: Methods and applications. J. Phys.: Condens. Matter, 18:R531–R583, 2006.

[7] Bai, X.C., McMullan, G. and Scheres, S.H. How cryo-EM is revolutionizing structural biology. Trends Biochem. Sci., 40:49–57, 2015.

[8] Lindorff-Larsen, K., Piana, S., Dror, R.O. and Shaw, D.E. How Fast-Folding Proteins Fold. Science, 334 (6055):517–520, 2011.

[9] Plattner, N., Doerr, S., Fabritiis, G.D. and Noé, F. Complete protein–protein association kinetics in atomic detail revealed by molecular dynamics simulations and Markov modelling. Nat. Chem., 9(10):1005, 2017.

[10] Wang, J. et al. Machine learning of coarse-grained molecular dynamics force fields. ACS Cent. Sci., 5:755–767, 2019.

[11] Charron, N.E. et al. Navigating protein landscapes with a machine-learned transferable coarse-grained model. 2310.18278, 2023.

[12] Noé, F., Olsson, S., Köhler, J. and Wu, H. Boltzmann Generators - Sampling equilibrium states of many-body systems with deep learning. Science, 365:eaaw1147, 2019.

[13] Zheng, S. et al. Predicting equilibrium distributions for molecular systems with deep learning. Nat. Mach. Intell., 6:558–567, 2024.

[14] Jing, B., Berger, B. and Jaakkola, T. AlphaFold meets flow matching for generating protein ensembles. 2402.04845, 2024.

[15] Qiao, Z., Nie, W., Vahdat, A., Iii, T.F.M. and Anandkumar, A. State-specific protein–ligand complex structure prediction with a multiscale deep generative model. Nat. Mach. Intell., 6:195–208, 2024.

[16] Wayment-Steele, H.K. et al. Predicting multiple conformations via sequence clustering and AlphaFold2. Nature, 625(7996):832–839, 2024.

[17] Bryant, P. and Noé, F. Structure prediction of alternative protein conformations. Nat. Commun., 15:7328, 2024.

[18] Vani, B.P., Aranganathan, A., Wang, D. and Tiwary, P. AlphaFold2-RAVE: From sequence to Boltzmann ranking. J. Chem. Theory Comput., 19:4351–4354, 2023.

[19] Aranganathan, A., Gu, X., Wang, D., Vani, B. and Tiwary, P. Modeling boltzmann weighted structural ensembles of proteins using ai based methods. ChemRxiv, 2024. doi: 10.26434/chemrxiv-2024-6f9h6-v2.

[20] Prinz, J.H. et al. Markov models of molecular kinetics: Generation and validation. J. Chem. Phys., 134:174105, 2011.

[21] Tsuboyama, K. et al. Mega-scale experimental analysis of protein folding stability in biology and design. Nature, 620(7973):434–444, 2023.

[22] Aviram, H.Y., Pirchi, M., Mazal, H. and Haran, G. Direct observation of ultrafast large-scale dynamics of an enzyme under turnover conditions. Proc. Natl. Acad. Sci. USA, 115:3243–3248, 2018.

[23] Fromowitz, F.B. et al. Ras p21 expression in the progression of breast cancer. Human pathology, 18(12):1268–1275, 1987.

[24] Greisman, J.B. et al. Discovery and validation of the binding poses of allosteric fragment hits to protein tyrosine phosphatase 1b: From molecular dynamics simulations to X-ray crystallography. J. Chem. Inf. Model., 63:2644–2650, 2023.

[25] Perez-Hernandez, G., Paul, F., Giorgino, T. D, Fabritiis, G. and Noé, F. Identification of slow molecular order parameters for markov model construction. J. Chem. Phys., 139:015102, 2013.

[26] Lane, T.J., Shukla, D., Beauchamp, K.A. and Pande, V.S. To milliseconds and beyond: Challenges in the simulation of protein folding. Curr. Opin. Struct. Biol., 23(1):58–65, 2013.

[27] Shaw, D.E. et al. Atomic-Level Characterization of the Structural Dynamics of Proteins. Science, 330 (6002):341–346, 2010.

[28] Chodera, J.D. and Noé, F. Markov state models of biomolecular conformational dynamics. Current Opinion in Structural Biology, 25:135–144, 2014.

[29] Best, R.B. and Mittal, J. Free-energy landscape of the gb1 hairpin in all-atom explicit solvent simulations with different force fields: Similarities and differences. Proteins, 79:1318–1328, 2011.

[30] Hahn, D.F., Gapsys, V., de Groot, B.L., Mobley, D.L. and Tresadern, G. Current state of open source force fields in protein-ligand binding affinity predictions. J. Chem. Inf. Model., 64:5063–5076, 2024.

[31] Laio, A. and Parrinello, M. Escaping free-energy minima. Proc. Natl. Acad. Sci. U.S.A., 99(20):12562– 12566, 2002.

[32] Robustelli, P., Piana, S. and Shaw, D.E. Developing a molecular dynamics force field for both folded and disordered protein states. Proc. Natl. Acad. Sci. U.S.A., 115(21), 2018.

[33] Sillitoe, I. et al. CATH: Increased structural coverage of functional space. Nucleic Acids Res., 49(D1): D266–D273, 2021.

[34] Malsam, J. et al. Complexin Suppresses Spontaneous Exocytosis by Capturing the Membrane-Proximal Regions of VAMP2 and SNAP25. Cell Reports, 32(3):107926, 2020.

[35] Zhou, Q. et al. The primed SNARE–complexin–synaptotagmin complex for neuronal exocytosis. Nature, 548(7668):420–425, 2017.

[36] Towler, P. et al. ACE2 X-Ray Structures Reveal a Large Hinge-bending Motion Important for Inhibitor Binding and Catalysis. J. Bio. Chem., 279(17):17996–18007, 2004.

[37] Zimmerman, M.I. et al. SARS-CoV-2 simulations go exascale to predict dramatic spike opening and cryptic pockets across the proteome. Nat. Chem., 13(7):651–659, 2021.

[38] Ouyang-Zhang, J., Diaz, D., Klivans, A. and Krähenbühl, P. Predicting a protein’s stability under a million mutations. Advances in Neural Information Processing Systems, 36:76229–76247, 2024.

[39] Cagiada, M., Ovchinnikov, S. and Lindorff-Larsen, K. Predicting absolute protein folding stability using generative models. bioRxiv, 2024. doi: 10.1101/2024.03.14.584940.

[40] Notin, P. et al. Proteingym: Large-scale benchmarks for protein fitness prediction and design. Advances in Neural Information Processing Systems, 36:64331–64379, 2024.

[41] Widatalla, T., Rafailov, R. and Hie, B. Aligning protein generative models with experimental fitness via direct preference optimization. bioRxiv, 2024. doi: 10.1101/2024.05.20.595026.

[42] Nikam, R., Kulandaisamy, A., Harini, K., Sharma, D. and Gromiha, M.M. ProThermDB: Thermodynamic database for proteins and mutants revisited after 15 years. Nucleic Acids Res., 49(D1):D420–D424, 2021.

[43] Tesei, G. et al. Conformational ensembles of the human intrinsically disordered proteome. Nature, 626 (8000):897–904, 2024.

[44] Tesei, G. and Lindorff-Larsen, K. Improved predictions of phase behaviour of intrinsically disordered proteins by tuning the interaction range. Open Research Europe, 2:94, 2023.

[45] Zhu, J. et al. Precise generation of conformational ensembles for intrinsically disordered proteins via fine-tuned diffusion models. bioRxiv, 2024. doi: 10.1101/2024.05.05.592611.

[46] Hofmann, H. et al. Polymer scaling laws of unfolded and intrinsically disordered proteins quantified with single-molecule spectroscopy. Proc. Natl. Acad. Sci. U.S.A., 109(40):16155–16160, 2012.

[47] Fröhlking, T., Bernetti, M., Calonaci, N. and Bussi, G. Toward empirical force fields that match experimental observables. J. Chem. Phys., 152:230902, 2020.

## References

[1] Steinegger, M. and Söding, J. MMseqs2 enables sensitive protein sequence searching for the analysis of massive data sets. Nature biotechnology, 35(11):1026–1028, 2017.

[2] van Kempen, M. et al. Fast and accurate protein structure search with Foldseek. Nat. Biotechnol., 42:243–246, 2024.

[3] Huguet, G. et al. Sequence-augmented SE (3)-flow matching for conditional protein backbone generation. 2405.20313, 2024.

[4] Krishna, R. et al. Generalized biomolecular modeling and design with RoseTTAFold All-Atom. Science, 384:eadl2528, 2024.

[5] Lindorff-Larsen, K., Piana, S., Dror, R.O. and Shaw, D.E. How fast-folding proteins fold. Science, 334 (6055):517–520, 2011.

[6] Hanson, S.M. et al. What Makes a Kinase Promiscuous for Inhibitors? Cell Chem. Biol., 26(3):390–399.e5, 2019.

[7] Chen, S. et al. The dynamic conformational landscape of the protein methyltransferase SETD8. eLife, 8:e45403, 2019.

[8] Thomson, E.C. et al. Circulating SARS-CoV-2 spike N439K variants maintain fitness while evading antibody-mediated immunity. Cell, 184(5):1171–1187.e20, 2021.

[9] Zimmerman, M.I. et al. SARS-CoV-2 simulations go exascale to predict dramatic spike opening and cryptic pockets across the proteome. Nat. Chem., 13(7):651–659, 2021.

[10] Abualrous, E.T. et al. MHC-II dynamics are maintained in HLA-DR allotypes to ensure catalyzed peptide exchange. Nat. Chem. Biol., 19(10):1196–1204, 2023.

[11] Plattner, N., Doerr, S., Fabritiis, G.D. and Noé, F. Complete protein–protein association kinetics in atomic detail revealed by molecular dynamics simulations and Markov modelling. Nat. Chem., 9(10):1005, 2017.

[12] Eastman, P. et al. OpenMM 7: Rapid development of high performance algorithms for molecular dynamics. PLOS Comput. Biol., 13(7):e1005659, 2017.

[13] Abraham, M.J. et al. GROMACS: High performance molecular simulations through multi-level parallelism from laptops to supercomputers. SoftwareX, 1–2:19–25, 2015.

[14] Jorgensen, W.L., Chandrasekhar, J., Madura, J.D., Impey, R.W. and Klein, M.L. Comparison of simple potential functions for simulating liquid water. J. Chem. Phys., 79(2):926–935, 1983.

[15] Hopkins, C.W., Le Grand, S., Walker, R.C. and Roitberg, A.E. Long-Time-Step Molecular Dynamics through Hydrogen Mass Repartitioning. J. Chem. Theory Comput., 11(4):1864–1874, 2015.

[16] Charron, N.E. et al. Navigating protein landscapes with a machine-learned transferable coarse-grained model. 2310.18278, 2023.

[17] Hruska, E., Abella, J.R., Nüske, F., Kavraki, L.E. and Clementi, C. Quantitative comparison of adaptive sampling methods for protein dynamics. The Journal of Chemical Physics, 149(24):244119, December 2018. ISSN 0021-9606, 1089-7690. doi: 10.1063/1.5053582.

[18] Scherer, M.K. et al. PyEMMA 2: A Software Package for Estimation, Validation, and Analysis of Markov Models. Journal of Chemical Theory and Computation, 11(11):5525–5542, November 2015. ISSN 1549-9618, 1549-9626. doi: 10.1021/acs.jctc.5b00743.

[19] Sillitoe, I. et al. CATH: Increased structural coverage of functional space. Nucleic Acids Research, 49(D1): D266–D273, 2021.

[20] Lindorff-Larsen, K. et al. Improved side-chain torsion potentials for the Amber ff99SB protein force field. Proteins, 78(8):1950–1958, 2010.

[22] Maier, J.A. et al. ff14SB: Improving the Accuracy of Protein Side Chain and Backbone Parameters from ff99SB. J. Chem. Theory Comput., 11(8):3696–3713, 2015.

[23] Robustelli, P., Piana, S. and Shaw, D.E. Developing a molecular dynamics force field for both folded and disordered protein states. Proc. Natl. Acad. Sci. U.S.A., 115(21), 2018.

[24] Piana, S., Lindorff-Larsen, K. and Shaw, D.E. How Robust Are Protein Folding Simulations with Respect to Force Field Parameterization? Biophys. J., 100(9):L47–L49, 2011.

[25] Duan, Y. et al. A point-charge force field for molecular mechanics simulations of proteins based on condensed-phase quantum mechanical calculations. J. Comput. Chem., 24(16):1999–2012, 2003.

[26] Hornak, V. et al. Comparison of multiple Amber force fields and development of improved protein backbone parameters. Proteins, 65(3):712–725, 2006.

[27] Jumper, J. et al. Highly accurate protein structure prediction with AlphaFold. Nature, 596(7873):583–589, 2021.

[28] Zheng, S. et al. Predicting equilibrium distributions for molecular systems with deep learning. Nat. Mach. Intell., 6:558–567, 2024.

[29] Mirdita, M. et al. Uniclust databases of clustered and deeply annotated protein sequences and alignments. Nucleic acids research, 45(D1):D170–D176, 2017.

[30] Remmert, M., Biegert, A., Hauser, A. and Söding, J. HHblits: Lightning-fast iterative protein sequence searching by HMM-HMM alignment. Nat. Meth., 9(2):173–175, 2012.

[31] Yim, J. et al. Se (3) diffusion model with application to protein backbone generation. 2302.02277, 2023.

[32] Ahdritz, G. et al. Openfold: Retraining AlphaFold2 yields new insights into its learning mechanisms and capacity for generalization. Nat. Meth., 21:1–11, 2024.

[33] Nichol, A. and Dhariwal, P. Improved denoising diffusion probabilistic models. 2102.09672, 2021.

[34] Bortoli, V.D. et al. Riemannian score-based generative modelling. 2202.02763, 2022.

[35] Song, Y. et al. Score-based generative modeling through stochastic differential equations. 2011.13456, 2020.

[36] Karras, T., Aittala, M., Aila, T. and Laine, S. Elucidating the design space of diffusion-based generative models. Advances in neural information processing systems, 35:26565–26577, 2022.

[37] Barrio-Hernandez, I. et al. Clustering predicted structures at the scale of the known protein universe. Nature, 622(7983):637–645, 2023.

[38] Prinz, J.H. et al. Markov models of molecular kinetics: Generation and validation. The Journal of Chemical Physics, 134(17):174105, 2011. ISSN 0021-9606, 1089-7690. doi: 10.1063/1.3565032.

[39] Wehmeyer, C. et al. Introduction to Markov state modeling with the PyEMMA software [Article v1.0]. LiveCoMS, 1(1):5965, 2018.

[40] Best, R.B., Hummer, G. and Eaton, W.A. Native contacts determine protein folding mechanisms in atomistic simulations. Proc. Natl. Acad. Sci. USA, 110(44):17874–17879, 2013.

[41] Luo, C. Understanding diffusion models: A unified perspective. arXiv preprint 2208.11970, 2022.

[42] Domingo-Enrich, C., Drozdzal, M., Karrer, B. and Chen, R.T. Adjoint matching: Fine-tuning flow and diffusion generative models with memoryless stochastic optimal control. 2409.08861, 2024.

[43] Xu, J. et al. Imagereward: Learning and evaluating human preferences for text-to-image generation. Advances in Neural Information Processing Systems, 36, 2024.

[44] Clark, K., Vicol, P., Swersky, K. and Fleet, D.J. Directly fine-tuning diffusion models on differentiable rewards. 2309.17400, 2023.

[45] Cimermancic, P. et al. CryptoSite: Expanding the druggable proteome by characterization and prediction of cryptic binding sites. J. Mol. Biol., 428(4):709–719, 2016.

[46] Meller, A., Bhakat, S., Solieva, S. and Bowman, G.R. Accelerating cryptic pocket discovery using alphafold. J. Chem. Theory Comput., 19(14):4355–4363, 2023.

[47] Chakravarty, D. and Porter, L.L. AlphaFold2 fails to predict protein fold switching. Prot. Sci., 31(6):e4353, 2022.

[48] Dana, J.M. et al. Sifts: updated structure integration with function, taxonomy and sequences resource allows 40-fold increase in coverage of structure-based annotations for proteins. Nucleic Acids Res., 47(D1): D482–D489, 2019.

[49] Jing, B., Berger, B. and Jaakkola, T. AlphaFold meets flow matching for generating protein ensembles. 2402.04845, 2024.

[50] Wayment-Steele, H.K. et al. Predicting multiple conformations via sequence clustering and AlphaFold2. Nature, 625(7996):832–839, 2024.

[51] Mirdita, M. et al. Colabfold: Making protein folding accessible to all. Nat. Meth., 19(6):679–682, 2022.

[52] Nikam, R., Kulandaisamy, A., Harini, K., Sharma, D. and Gromiha, M.M. ProThermDB: Thermodynamic database for proteins and mutants revisited after 15 years. Nucleic Acids Res., 49(D1):D420–D424, 2021.

[53] Tesei, G. et al. Conformational ensembles of the human intrinsically disordered proteome. Nature, 626 (8000):897–904, 2024.

[54] Zhu, J. et al. Precise generation of conformational ensembles for intrinsically disordered proteins via fine-tuned diffusion models. bioRxiv, 2024. doi: 10.1101/2024.05.05.592611.

[55] Pérez-Hernández, G., Paul, F., Giorgino, T., Fabritiis, G.D. and Noé, F. Identification of slow molecular order parameters for Markov model construction. J. Chem. Phys., 139(1):015102, 2013.

[56] Hoffmann, M. et al. Deeptime: A Python library for machine learning dynamical models from time series data. Mach. Learn.: Sci. Technol., 3(1):015009, 2022.

